# miR-324 mediates bone homeostasis through the regulation of osteoblast and osteoclast differentiation and activity

**DOI:** 10.1101/2023.07.10.548366

**Authors:** Dan J. Hayman, Hua Lin, Amanda Prior, Gemma Charlesworth, Francesca M. Johnson de Sousa Brito, Yao Hao, Krutik Patel, Jamie Soul, Ian M. Clark, Katarzyna A. Piróg, Matt J. Barter, Rob Van ’T Hof, David A. Young

## Abstract

microRNAs (miRNAs) are non-coding RNAs which modulate the expression of other RNA molecules. One miRNA can target many transcripts, allowing each miRNA to play key roles in many biological pathways. miR-324 is a miRNA previously implicated in bone and cartilage maintenance, defects of which result in common age-related diseases, such as osteoporosis or osteoarthritis (OA).

In global miR-324-null mice cartilage damage was increased in both surgically and ageing-induced OA, despite minimal changes to the cartilage transcriptome, with few predicted miR-324 targets dysregulated. However, micro-computed tomography and histology demonstrated that global miR- 324-null the mice had an increase in bone mineral density, trabecular thickness and cortical thickness, with many parameters increasing with age. The bone marrow of miR-324-null mice also had reduced lipid content while and *in vivo* TRAP staining revealed a decrease in osteoclasts, with histomorphometry demonstrating an increased rate of bone formation in miR-324-null mice.

*Ex vivo* assays revealed that the high bone mass phenotype of the miR-324-null mice resulted from increased osteoblast activity and decreased osteoclastogenesis. RNA-seq and qRT-PCR followed by miR-324 target prediction and validation in osteoblasts, osteoclasts and bone marrow macrophages identified the osteoclast fusion regulator *Pin1* as a miR-324 target in the osteoclast lineage and the master osteogenic regulator *Runx2* as a target of miR-324-5p in osteoblasts, the *in vitro* overexpression of which recapitulated the increased osteogenesis and decreased adipogenesis phenotype observed *in vivo*.

These data point to important roles of miR-324 in skeletal biology with altered bone homeostasis in miR-324-null mice potentially causal for the increased cartilage damage observed during OA and ageing. Elucidation of pathways regulated by miR-324 offer promise for the treatment of bone diseases such as osteoporosis.

## Introduction

microRNAs (miRNAs) are a class of non-coding RNA which commonly modulate the expression of target RNA molecules through complimentary binding to the 3’ untranslated region (3’UTR) of target mRNAs. As a consequence, the mRNA target is generally degraded. One miRNA can target many mRNAs and each mRNA is often the target of multiple miRNAs (1).

With an ageing demographic, musculoskeletal conditions are increasing in prevalence. The most common musculoskeletal conditions are osteoarthritis (OA) and osteoporosis (OP) (2, 3). Treatments for OA are limited to analgesic use prior to eventual joint replacement. For OP therapeutic options are better and range from supplements (e.g., calcium and vitamin D), through to hormone replacement, bisphosphonates and more recently biologics such as Denosumab and Romosozumab, amongst other treatments (3). Studies have suggested that OA and OP are inversely related, depending on the definition of OA (4). During OA progression, in addition to cartilage degradation, bone remodelling occurs. At late stages of the condition the subchondral bone of OA patients is severely thickened due to dysregulation of osteoblasts and osteoclasts, the cells responsible for bone formation and resorption, respectively (5, 6)

Bone remodelling is a crucial process, responsible for the maintenance of bone mass (7). It is comprised of antagonistic effects provided by osteoblasts and osteoclasts, along with osteocytes, which regulate the former cells (8). The differentiation of these cells from mesenchymal and haematopoietic stem cells in the bone marrow is cross-regulated by mature bone cells, such that bone remodelling can be precisely modulated according to the need of the skeletal system at a particular time. miRNAs play numerous roles in the processes involved in bone remodelling. To date, more than 80 miRNAs have been reported to affect either osteogenesis, osteoclastogenesis or the mediation of these processes by osteocytes (9). For example, the miR-185-null mouse exhibits dramatically increased bone formation due to increased osteogenesis (10), whereas miR-124 targets NFATc1, the transcription factor responsible for activation of osteoclastogenic genes, thus acting to repress osteoclastogenesis (11). Due to the complex networks through which these processes communicate, a single miRNA can therefore have large implications on the underlying balance between bone formation and resorption.

miR-324 is a miRNA which has been associated with diseases of the cartilage and bone, including bone formation and as a serum marker for fracture risk (12–15). miR-324 can reportedly regulate aspects of the Hedgehog (Hh) signalling (15–17), a pathway instrumental for many musculoskeletal processes including endochondral ossification, whereby cartilage tissue is replaced by bone (18). Here we demonstrate that in a global knockout mouse model of *Mir324*, cartilage damage is increased both spontaneously and following surgical induction of OA, despite harbouring few dysregulated genes in chondrocytes. However, the miR-324-null mice also display a high bone mass phenotype, resulting from a decrease in osteoclastogenesis and an increase in bone formation.

Finally, we functionally define bone development and disease-relevant targets of miR-324.

## Results

### miR-324-null mice display increased cartilage damage

miR-324-5p has previously been associated with human OA (15, 19) while serum levels of miR-324- 3p show correlation with bone mineral density (BMD), mineral apposition rate (MAR), bone formation rate (BFR) and mineralised bone surface (12–14). However, miR-324-related musculoskeletal phenotypes have until now remained unexplored *in vivo*. Having previously generated a miR-324-null mouse (20), these and WT counterparts were subjected to trauma-induced OA using the destabilisation of the medial meniscus (DMM) model (21). miR-324-null mice displayed a significant increase in cartilage damage at the medial femoral condyle (MFC), the medial tibial plateau (MTP) and with the summed scores at 5.3-months of age (8-weeks post-surgery). miR-324- null mice aged to 14-months also showed increased spontaneous cartilage damage (Figures 1a and 1b).

**Figure 1.**
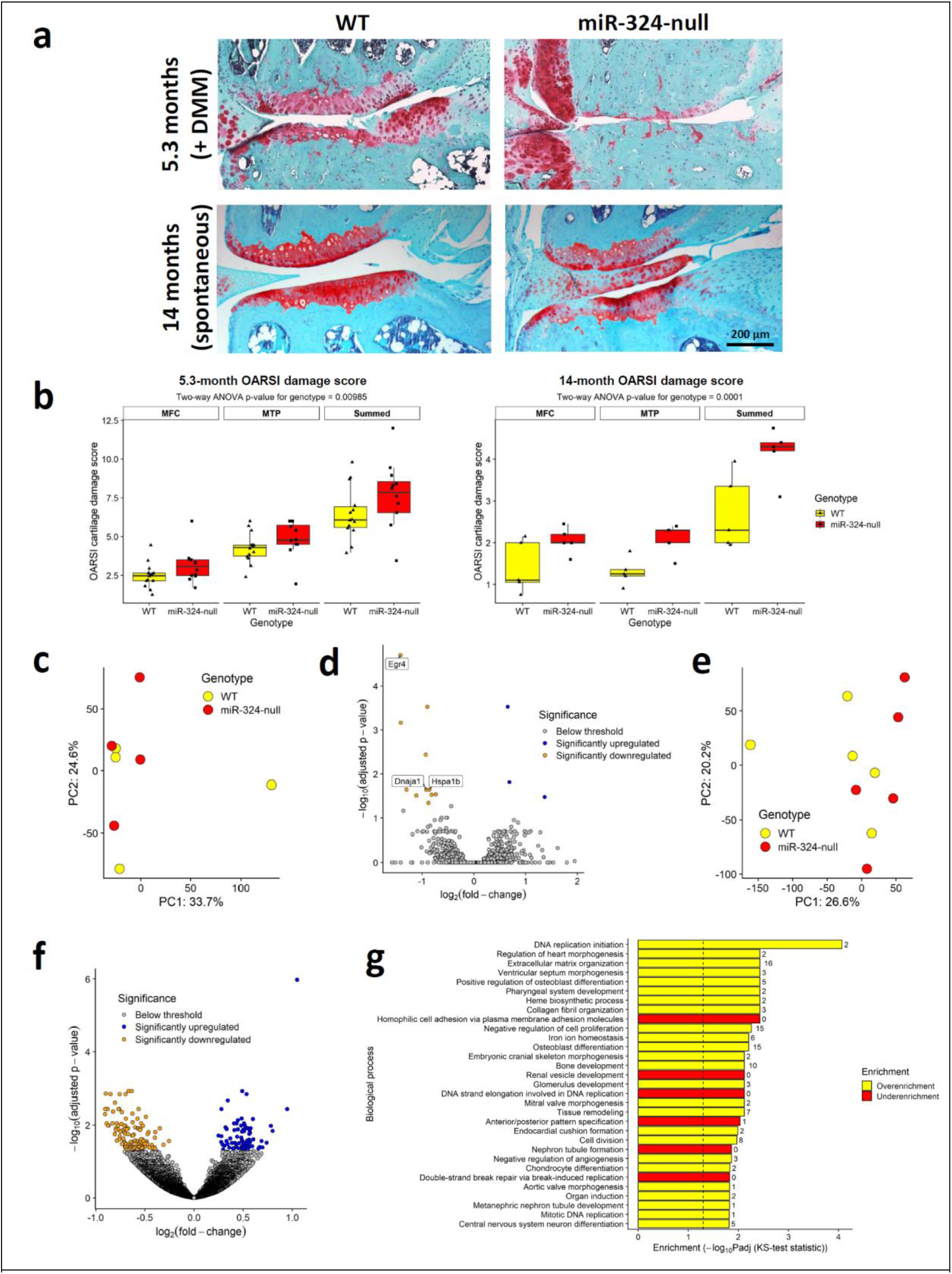
miR-324-null mice display an increase in cartilage damage despite limited differential expression in the chondrocyte transcriptome. Cartilage damage was assessed in two cohorts of miR-324-null and WT mice. To induce post-traumatic OA, miR-324-null and WT mice aged 16- weeks underwent DMM surgery and were subsequently sacrificed at 24-weeks of age (5.3- months; N = 14 WT and 12 miR-324-null mice). To investigate whether miR-324 affects spontaneous cartilage damage, miR-324-null and WT mice were aged for 14-months, before being sacrificed (N = 5 mice per genotype). For both cohorts, after sacrifice the left hind leg processed, sectioned and subsequently stained with Safranin-O to assess cartilage damage the using the OARSI histopathological scoring system (22). (**a**) Representative images of the medial condyle of miR-324-null and WT mice in each condition. In both spontaneous and DMM-induced cohorts, cartilage damage is worsened by the lack of miR-324. (**b**) Quantification of cartilage damage confirms that damage is worsened in miR-324-null mice relative to WT controls. P-values for genotype were determined using two-way ANOVA across the two joint regions (MFC, medial femoral condyle and MTP, medial tibial plateau) and overall summed score. (**c**) RNA was extracted and sequenced from the costal chondrocytes of miR-324-null and WT mice aged 7-days. Samples did not segregate well by PCA. (**d**) 15 genes were significantly differentially expressed between the two genotypes (adjusted p-value ≤ 0.05 and TPM ≥ 5), of which predicted miR-324 targets are labelled. (e) Knee cartilage was dissected from 5 miR-324-null and 5 WT mice aged 20-weeks, and the RNA extracted for sequencing. Samples did not segregate well by PCA. (**f**) 198 genes were differentially expressed (adjusted p-value ≤ 0.05 and WT TPM ≥ 5) between miR-324-null and WT cartilage. (**g**) GO term enrichment of the knee cartilage RNA samples, revealed several cartilage-related GO terms as significantly enriched, however a higher number of enriched GO terms related to bone. The dotted line indicates the significance threshold used to identify statically significantly enriched terms (adjusted p-value . 0.05, using fold-change ranks as the input), utilising a Kolmogorov-Smirnov (KS) test, and the number of statistically significant genes in each term is shown to the right-hand side of the bar.

To explain this increased cartilage damage phenotype in miR-324-null mice, we undertook an RNA- seq experiment between miR-324-null and WT chondrocytes, isolated form costal growth plates. The samples did not segregate by genotype using principal component analysis (PCA) (Figure 1c and Supplemental data table 1) and in total only 15 genes were significantly differentially expressed between the two genotypes (adjusted p-value ≤ 0.05 and TPM ≥ 5), in spite of the increased cartilage damage observed (Figure 1d). Furthermore, only 3 significantly differentially expressed genes were predicted targets of miR-324 according to TargetScan (*Dnaja1*, *Egr4* and *Hspa1b*; labelled in Figure 1c), and all 3 of these genes were downregulated in the miR-324-null samples, suggesting that these are not direct miR-324 target genes in chondrocytes and are therefore unlikely to be responsible for the increased cartilage damage observed. In addition, no Gene Ontology (GO) terms were significantly enriched in these data (data not shown). To confirm that genetic dysregulation in cartilage did not arise with age in miR-324-null mice, we sequenced RNA from the tibial medial condyle of knee cartilage of healthy 20-week-old miR-324-null and WT mice. This dissected tissue is composed primarily of cartilage but also contains the underlying subchondral bone. An increased number of genes were differentially expressed between the two genotypes (198 genes with adjusted p-value ≤ 0.05 and WT TPM ≥ 5, consisting of 28 significantly upregulated predicted miR-324 targets; Figure 1e and f and Supplemental data table 2) and 3 matrix-related GO terms were significantly enriched (Extracellular matrix organisation, Collagen fibril organisation and Chondrocyte differentiation). However, a greater number of enriched GO terms related to bone (Figure 1g). This evidence led us to hypothesise that the increased cartilage damage in miR-324-null mice may not exclusively be due to altered cartilage maintenance pathways, but rather an indirect consequence of dysregulation in bone. This hypothesis was supported by the increased abundance of miR-324 in bone cells relative to cartilage (Supplementary Figure S1).

### miR-324-null mice display a high bone mass phenotype

Through the utilisation of micro-computed tomography (µCT), we assessed whether the bone microstructure of male miR-324-null mice differed from that of male wild-type mice (WTs), aged 5-, 7- and 14-months. Although miR-324-null mice lacked any obvious alterations to limb development or body size (data not shown), they did display increased bone thickness, both in the femoral and tibial trabeculae and cortex. Although trabecular bone thickness remained increased in miR-324-null samples relative to WT controls, even at 14-months, the increase in miR-324-null cortical thickness reduced with age (Figure 2 and Tables 1).

**Figure 2.**
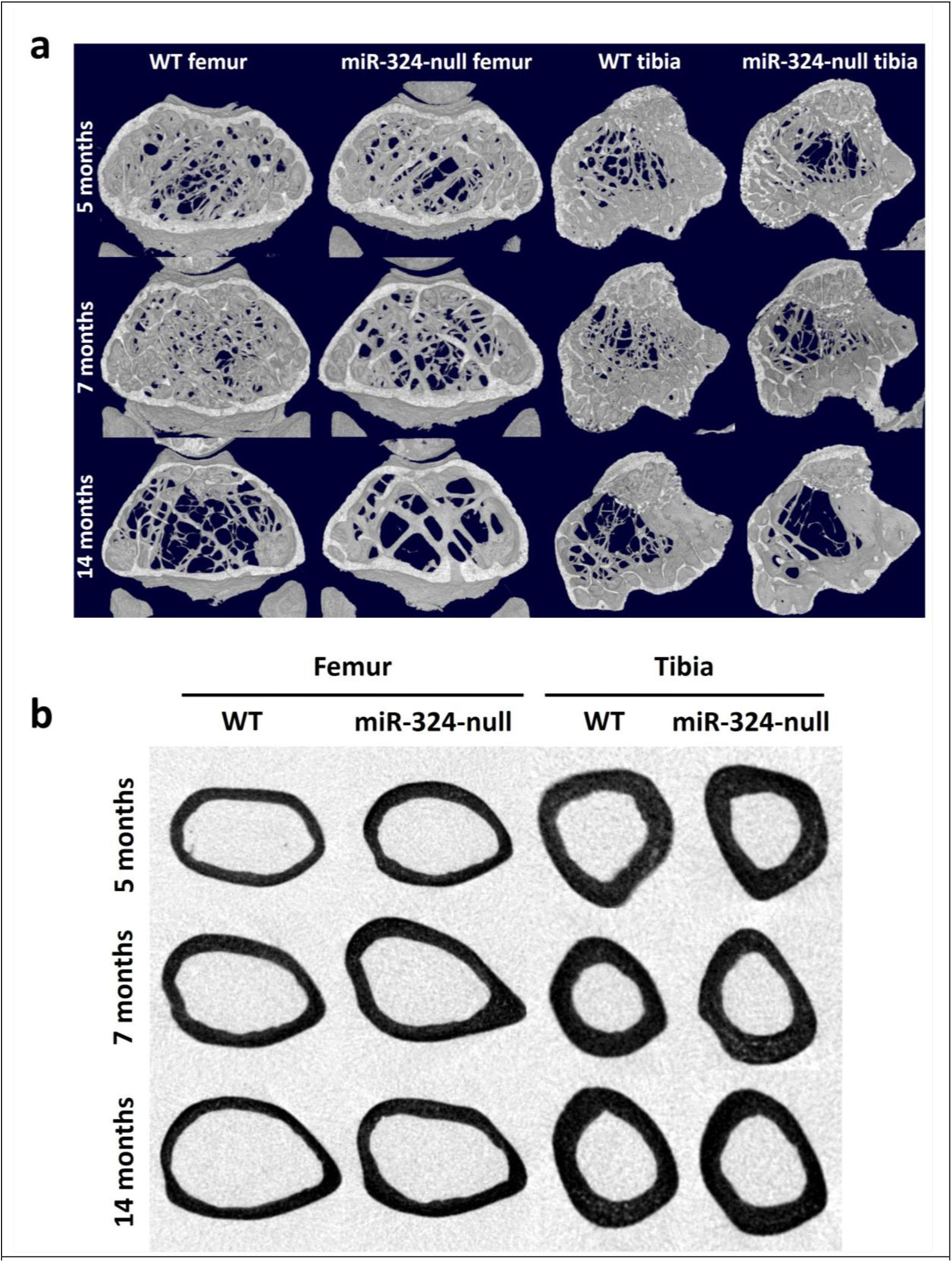
Representative images of the miR-324-null increased bone mass phenotype. (**a**) Representative trabecular femoral and tibial µCT reconstructions are shown for male miR-324-null and WT mice aged 5-, 7- and 14-months. (**b**) Representative slices on the transaxial plane, taken from the midpoint of either the tibia or femur, are shown. Increased thickness can be observed in both femoral and tibial miR-324-null cortices, relative to WT age-matched controls, across the time points measured.

In addition to the miR-324-null increased bone thickness phenotype, the trabecular structural model index (SMI) was also increased in both femoral and tibial miR-324-null bones. The overall effect of genotype on SMI was statistically significant for both the femur and tibia, suggestive of more rod-like trabeculae in the miR-324-null mice and therefore a less mechanically competent structure (23). SMI increased with age in both genotypes of our mice, as previously documented in humans (24), and therefore the miR-324-null mice may display an accelerated ageing phenotype. The bone tissue mineral density (TMD) of miR-324-null mice was also increased relative to WT mice in both the femur and tibia (Table 1). In both trabecular and cortical bone this increase was most apparent at the 5-month time point in cortical bone but decreased with age, with no observable difference by the 14-month time point. Due to the observed early increase in TMD, we also investigated whether any bowing could be observed in the tibiae of miR-324-null mice aged 5-, 7- and 14-months, by measuring the distance between the femur and tibia at the vertical tibial midpoint. A greater horizontal distance between the fibula and tibia at the tibial midpoint was observed, an increase of approximately 5%, showing that miR-324-null tibiae have increased bowing relative to WT controls (Supplementary Figure S2).

**Table 1.**
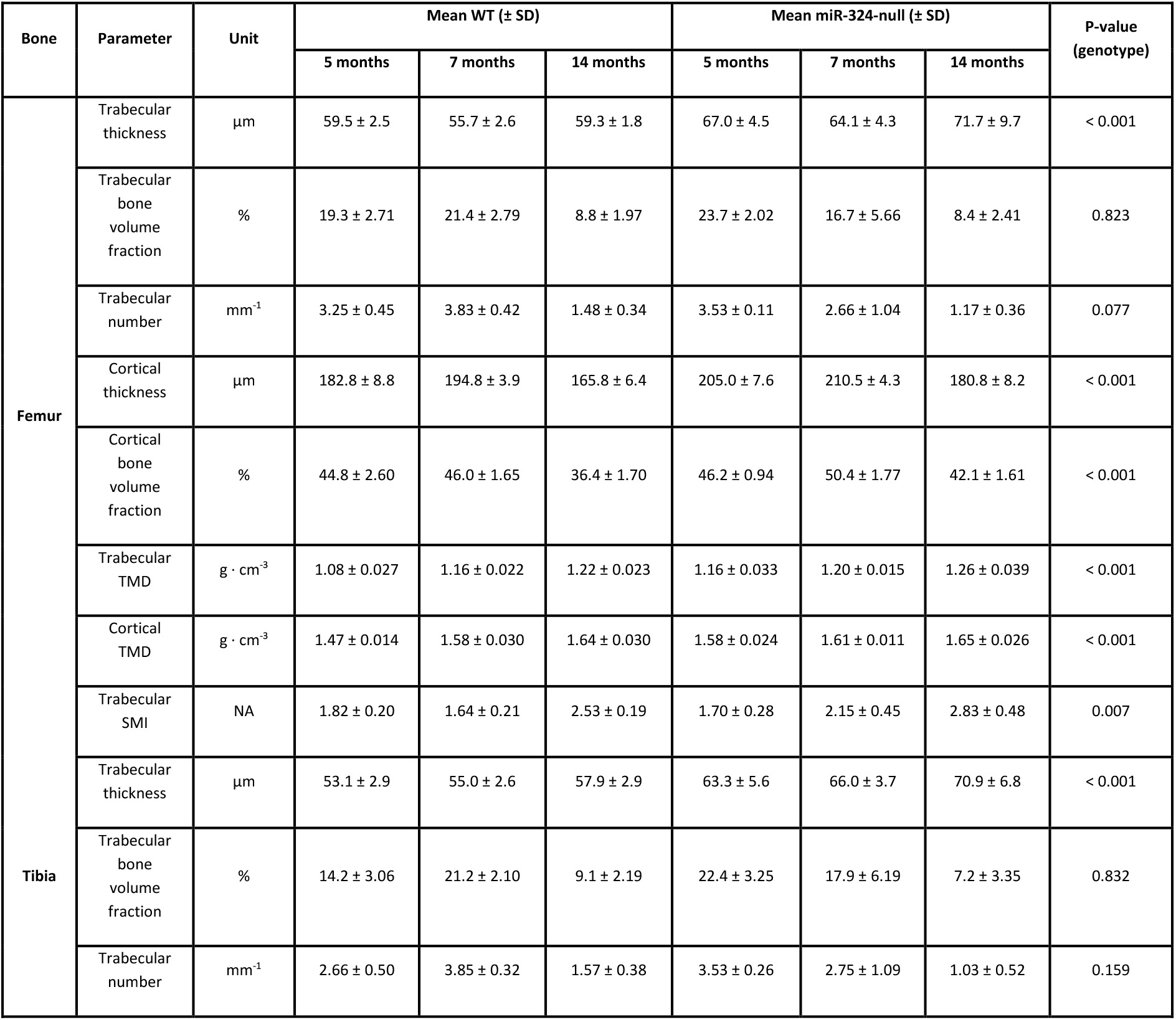
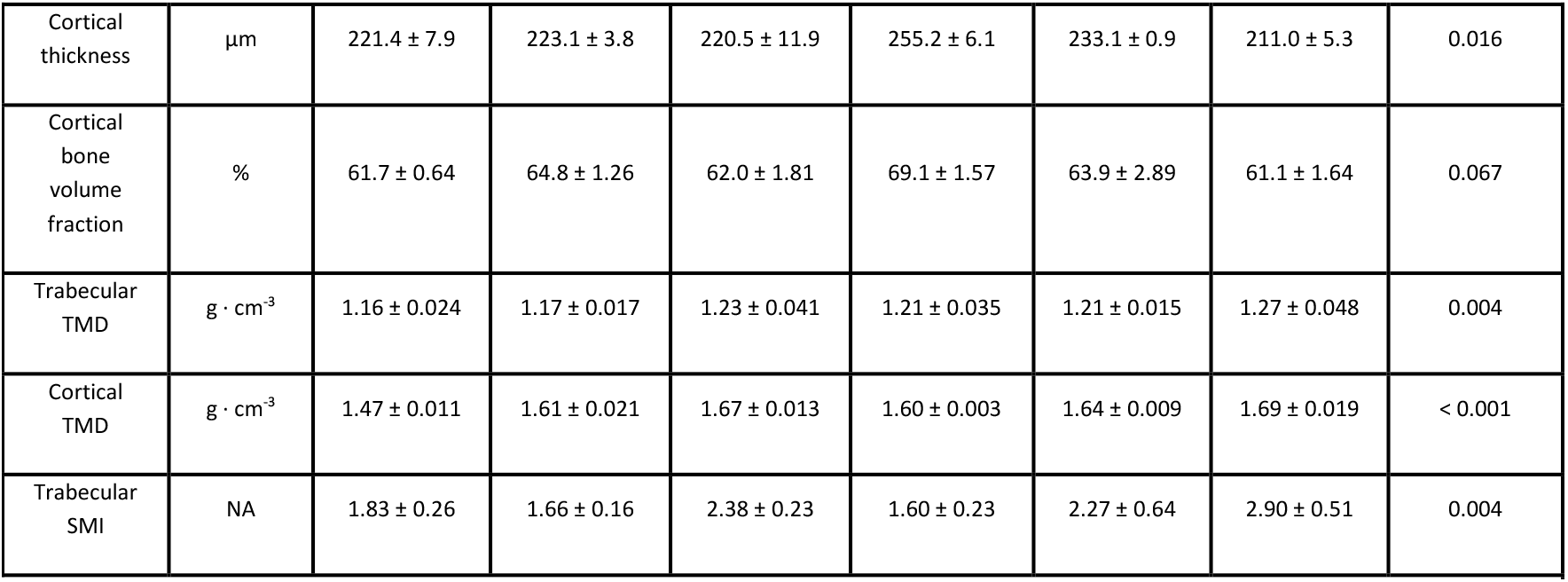
miR-324-null mice display altered femoral and tibial bone parameters. Each parameter was measured using µCT. The mean value ± SD for each parameter, genotype and age is shown. P-values for genotype were calculated using ANOVA. NA, not applicable.

| Bone | Parameter | Unit | Mean WT (± SD) | | | Mean miR-324-null (± SD) | | | P-value (genotype) |
| --- | --- | --- | --- | --- | --- | --- | --- | --- | --- |
|  |  |  | 5 months | 7 months | 14 months | 5 months | 7 months | 14 months |  |
| Femur | Trabecular thickness | $\mu\text{m}$ | 59.5 ± 2.5 | 55.7 ± 2.6 | 59.3 ± 1.8 | 67.0 ± 4.5 | 64.1 ± 4.3 | 71.7 ± 9.7 | < 0.001 |
| | Trabecular bone volume fraction | % | 19.3 ± 2.71 | 21.4 ± 2.79 | 8.8 ± 1.97 | 23.7 ± 2.02 | 16.7 ± 5.66 | 8.4 ± 2.41 | 0.823 |
| | Trabecular number | $\text{mm}^{-1}$ | 3.25 ± 0.45 | 3.83 ± 0.42 | 1.48 ± 0.34 | 3.53 ± 0.11 | 2.66 ± 1.04 | 1.17 ± 0.36 | 0.077 |
| | Cortical thickness | $\mu\text{m}$ | 182.8 ± 8.8 | 194.8 ± 3.9 | 165.8 ± 6.4 | 205.0 ± 7.6 | 210.5 ± 4.3 | 180.8 ± 8.2 | < 0.001 |
| | Cortical bone volume fraction | % | 44.8 ± 2.60 | 46.0 ± 1.65 | 36.4 ± 1.70 | 46.2 ± 0.94 | 50.4 ± 1.77 | 42.1 ± 1.61 | < 0.001 |
| | Trabecular TMD | $\text{g} \cdot \text{cm}^{-3}$ | 1.08 ± 0.027 | 1.16 ± 0.022 | 1.22 ± 0.023 | 1.16 ± 0.033 | 1.20 ± 0.015 | 1.26 ± 0.039 | < 0.001 |
| | Cortical TMD | $\text{g} \cdot \text{cm}^{-3}$ | 1.47 ± 0.014 | 1.58 ± 0.030 | 1.64 ± 0.030 | 1.58 ± 0.024 | 1.61 ± 0.011 | 1.65 ± 0.026 | < 0.001 |
| | Trabecular SMI | NA | 1.82 ± 0.20 | 1.64 ± 0.21 | 2.53 ± 0.19 | 1.70 ± 0.28 | 2.15 ± 0.45 | 2.83 ± 0.48 | 0.007 |
| Tibia | Trabecular thickness | $\mu\text{m}$ | 53.1 ± 2.9 | 55.0 ± 2.6 | 57.9 ± 2.9 | 63.3 ± 5.6 | 66.0 ± 3.7 | 70.9 ± 6.8 | < 0.001 |
| | Trabecular bone volume fraction | % | 14.2 ± 3.06 | 21.2 ± 2.10 | 9.1 ± 2.19 | 22.4 ± 3.25 | 17.9 ± 6.19 | 7.2 ± 3.35 | 0.832 |
| | Trabecular number | $\text{mm}^{-1}$ | 2.66 ± 0.50 | 3.85 ± 0.32 | 1.57 ± 0.38 | 3.53 ± 0.26 | 2.75 ± 1.09 | 1.03 ± 0.52 | 0.159 |

|  |  |  |  |  |  |  |  |  |
| --- | --- | --- | --- | --- | --- | --- | --- | --- |
| Cortical thickness | μm | 221.4 ± 7.9 | 223.1 ± 3.8 | 220.5 ± 11.9 | 255.2 ± 6.1 | 233.1 ± 0.9 | 211.0 ± 5.3 | 0.016 |
| Cortical bone volume fraction | % | 61.7 ± 0.64 | 64.8 ± 1.26 | 62.0 ± 1.81 | 69.1 ± 1.57 | 63.9 ± 2.89 | 61.1 ± 1.64 | 0.067 |
| Trabecular TMD | g · cm<sup>-3</sup> | 1.16 ± 0.024 | 1.17 ± 0.017 | 1.23 ± 0.041 | 1.21 ± 0.035 | 1.21 ± 0.015 | 1.27 ± 0.048 | 0.004 |
| Cortical TMD | g · cm<sup>-3</sup> | 1.47 ± 0.011 | 1.61 ± 0.021 | 1.67 ± 0.013 | 1.60 ± 0.003 | 1.64 ± 0.009 | 1.69 ± 0.019 | < 0.001 |
| Trabecular SMI | NA | 1.83 ± 0.26 | 1.66 ± 0.16 | 2.38 ± 0.23 | 1.60 ± 0.23 | 2.27 ± 0.64 | 2.90 ± 0.51 | 0.004 |

Considering the high bone mass phenotype observed in miR-324-null mice, we proceeded to investigate whether this could be explained by increased osteoblast-mediated bone formation. To assess the rate of bone formation for the differing between genotypes, 7- and 14-month-old mice were injected with alizarin red-S followed by calcein (Table 2). This revealed an increased mineral apposition rate (MAR) and area of mineralising surface in 14-month miR-324-null mice, as well as a significantly increased bone formation rate (BFR) at both 7- and 14-months. Additionally, Goldner’s Trichrome staining revealed that miR-324-null mice had thickened osteoid, the unmineralised component of bone, at both ages. The miR-324-null tibial sections also displayed a reduction in lipid droplets, both in terms of their number and mean area, although lipid droplet area reduction was not statistically significant in the 7-month cohort (Table 2 and Figure 3). Combined, these results suggest that lack of *Mir324* in mice may result in the observed high bone mass phenotype at least in part through modulating the balance between osteogenesis and adipogenesis.

**Table 2.** Histomorphology and histological staining reveal that miR-324-null mice display increased osteoblast-mediated bone formation and reduced leg bone marrow lipid content. Formalin-fixed tibial bone sections from miR-324-null and WT mice aged 7- or 14-months were stained either with Goldner’s Trichrome (for osteoid) or alizarin red-S and calcein green (for bone formation parameters) and analysed using OsteoidHisto and CalceinHisto (25). The mean value ± SD for each parameter, genotype and age is shown. P-values for genotype were calculated using ANOVA.

| Parameter | Unit | Mean WT (± SD) |  | Mean miR-324-null (± SD) |  | P-value (genotype) |
| --- | --- | --- | --- | --- | --- | --- |
|  |  | 7 months | 14 months | 7 months | 14 months |  |
| Mineralising surface | % | 15.6 ± 3.4 | 15.2 ± 2.9 | 24.7 ± 10.5 | 19.3 ± 3.5 | 0.027 |
| MAR | μm day<sup>-1</sup> | 0.23 ± 0.096 | 0.26 ± 0.026 | 0.32 ± 0.030 | 0.40 ± 0.098 | 0.005 |
| BFR | μm day<sup>-1</sup> | 0.036 ± 0.019 | 0.040 ± 0.006 | 0.092 ± 0.020 | 0.062 ± 0.010 | < 0.001 |
| Mean osteoid thickness | μm | 0.60 ± 0.045 | 0.65 ± 0.010 | 0.69 ± 0.021 | 0.71 ± 0.020 | < 0.001 |
| Osteoid surface per bone surface | % | 3.68 ± 1.4 | 55.55 ± 23.6 | 10.68 ± 4.4 | 69.13 ± 22.4 | 0.144 |
| Lipid droplets per tissue area | nm<sup>-2</sup> | 0.91 ± 0.24 | 4.75 ± 2.35 | 0.30 ± 0.08 | 0.34 ± 0.23 | 0.001 |
| Mean area of each lipid droplet | $\mu\text{m}^2$ | $55.9 \pm 30.82$ | $93.5 \pm 7.61$ | $22.5 \pm 3.53$ | $62.0 \pm 20.43$ | 0.008 |

**Figure 3.**
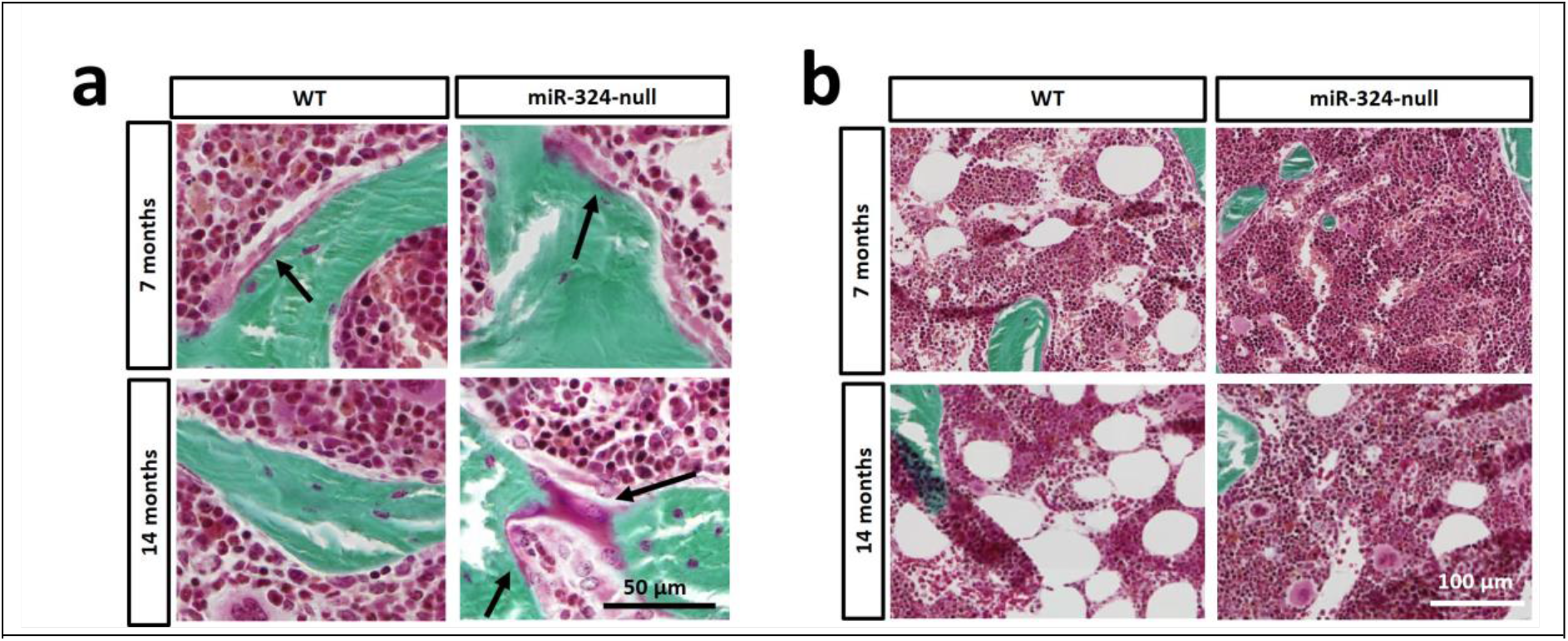
miR-324-null mice display altered bone and lipid formation *in vivo*. Formalin-fixed tibial bone sections from miR-324-null and WT mice aged 7- or 14-months were stained with Goldner’s Trichrome, which labels mineralised bone green, in comparison to the osteoid (unmineralised bone), which is labelled pink. (**a**) Increased osteoid is observed in miR-324-null

### miR-324-null osteoblasts display increased bone formation *ex vivo*

Since *in vivo* miR-324-null samples displayed evidence of increased bone formation, we next examined the phenotype of isolated and cultured murine calvariae pre-osteoblasts *ex vivo*. Isolated cells were cultured until confluent (14 days), stimulated with osteogenic media (for 7 days), before being either stained with alizarin red-S or assessed for alkaline phosphatase activity. miR-324-null osteoblasts showed a statistically significant increase of approximately 5% in alizarin red-S intensity and an increase of approximately 30% in alkaline phosphatase activity. Gene expression was analysed in parallel cultures, with the expression of the key osteogenesis-related genes *Alpl*, *Col1a1*, *Col1a2* and *Runx2* significantly upregulated in miR-324-null osteoblasts (Figure 4a). To confirm that the alterations in *ex vivo* mineralisation and gene expression were a consequence of miR-324 removal, both arms of the miRNA were reintroduced into miR-324-null osteoblasts, at initially determined endogenous levels (Supplementary Figure S3). Upon transfection with the mimics the increased alizarin red-S staining previously observed in miR-324-null osteoblasts was significantly reduced. Concomitantly, the expression of *Col1a2* and *Runx2* were significantly reduced, whilst *Col1a1* expression was also lower but not-significantly (p-value = 0.054). Interestingly, addition of the miR-324 mimics did not alter *Alpl* expression, implying that this miR-324-null-driven dysregulation cannot be reversed by transient reintroduction of miR-324-5p and -3p (Figures 4bi and 4bii). Runx2 protein levels were also returned to physiological levels with the re-introduction of miR- 324 (Figure 4biii).

**Figure 4.**
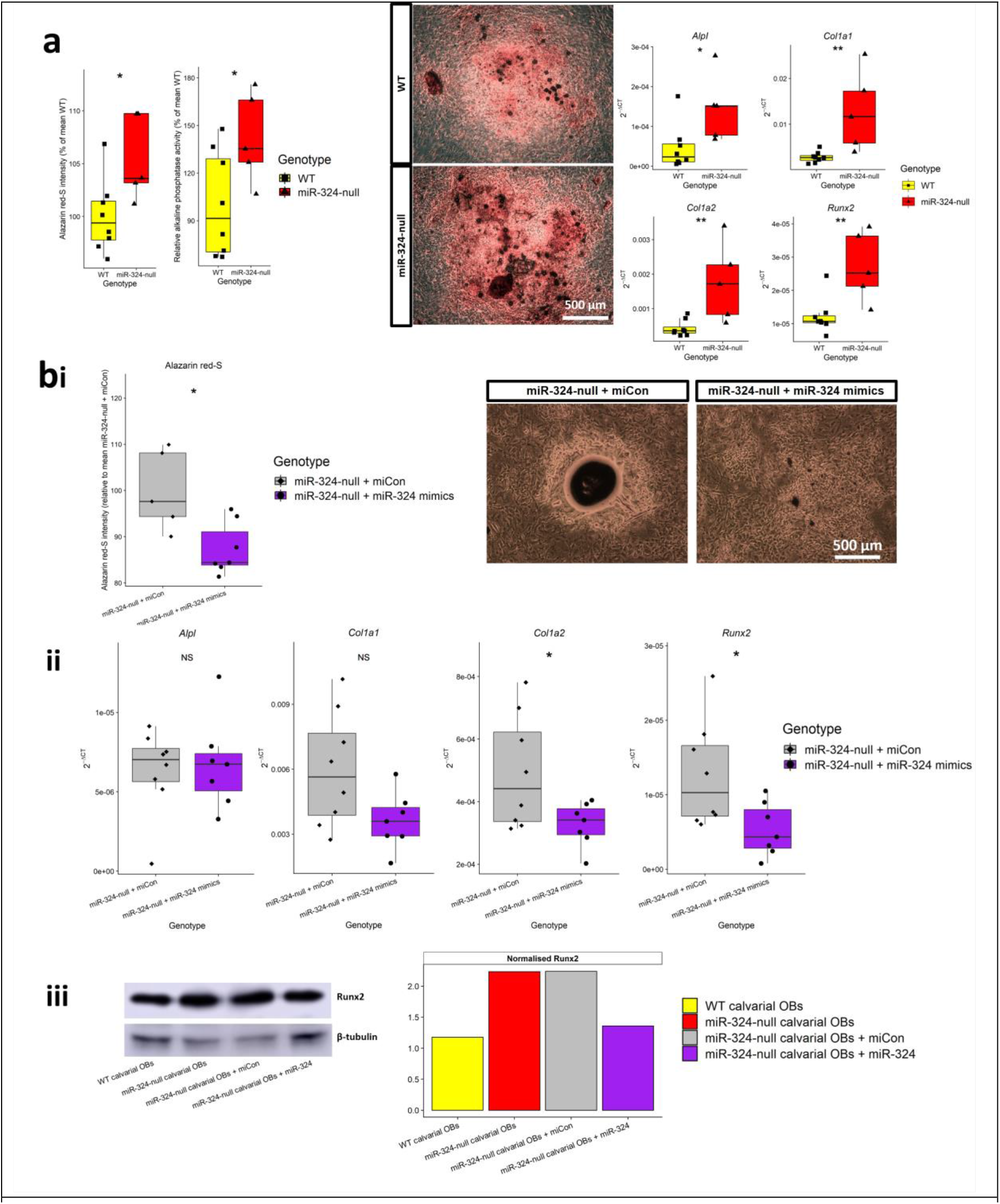
miR-324-null osteoblasts display increased bone formation, which can be rescued by the re-introduction of physiological concentrations of miR-324. (**a**) Calvarial osteoblasts were isolated from miR-324-null and WT pups aged 3-days and cultured in osteogenic media for 7-days, after which parallel cultures were stained for alizarin red-S, assessed for alkaline phosphatase activity, or RNA extracted, and expression of key bone formation marker genes assessed. miR- 324-null calvarial osteoblasts displayed increased bone formation by all of these parameters. (**b**) miR-324-null calvarial osteoblasts were transfected either with miCon (negative control miRNA mimic) or miR-324-5p and -3p mimics at physiological concentrations (Supplementary Figure S3) for 24 hours. (**i**) Osteoblasts transfected with miR-324 mimics displayed reduced bone formation, determined by alizarin red-S staining following 7-days of culture in osteogenic media. (**ii**) Transfection with miR-324 mimics at physiological concentrations also repressed *Col1a2* and *Runx2* expression, quantified using RT-qPCR. (**iii**) Transfection with miR-324 mimics at physiological concentrations rescued the excessive Runx2 levels in pooled miR-324-null osteoblasts isolated from 5 mice. The treatments were as follows: WT, miR-324-null, miR-324-null + 50nM miCon and miR-324-null + 50nM miR-324 mimics. All groups were treated with osteogenic media for 7-days. For all RT-qPCR experiments, 18S was utilised as the housekeeping gene. For all panels, ** and * represent p-values ≤ 0.01 and 0.05, respectively, calculated using two-tailed Student’s *t*-tests. Each point represents osteoblast cultures isolated from a single mouse.

To identify the origin of the abnormal bone formation in the osteoblasts of miR-324-null mice, osteoblasts were isolated from hind leg bone chips of 20-week-old miR-324-null and WT mice and grown to confluency for approximately 6 weeks, before being stimulated with osteogenic media for 18 days. RNA-seq was subsequently undertaken, revealing that more than 3500 genes were significantly differentially expressed (adjusted p-value ≤ 0.05) between miR-324-null and WT osteoblasts (Figure 5a and Supplemental data table 3), with PCA revealing clear segregation of the samples by genotype (Figure 5b). As observed in the *ex vivo* calvarial osteoblasts, osteogenic genes *Col1a1* and *Runx2* were within the significantly upregulated genes. No obvious Gene Ontology pathways related to bone formation were enriched in the top 20 pathways (Figure 5c), although “Arthritis” and “Rheumatoid arthritis” were enriched in Disease Ontology analysis (Figure 5d).

**Figure 5.**
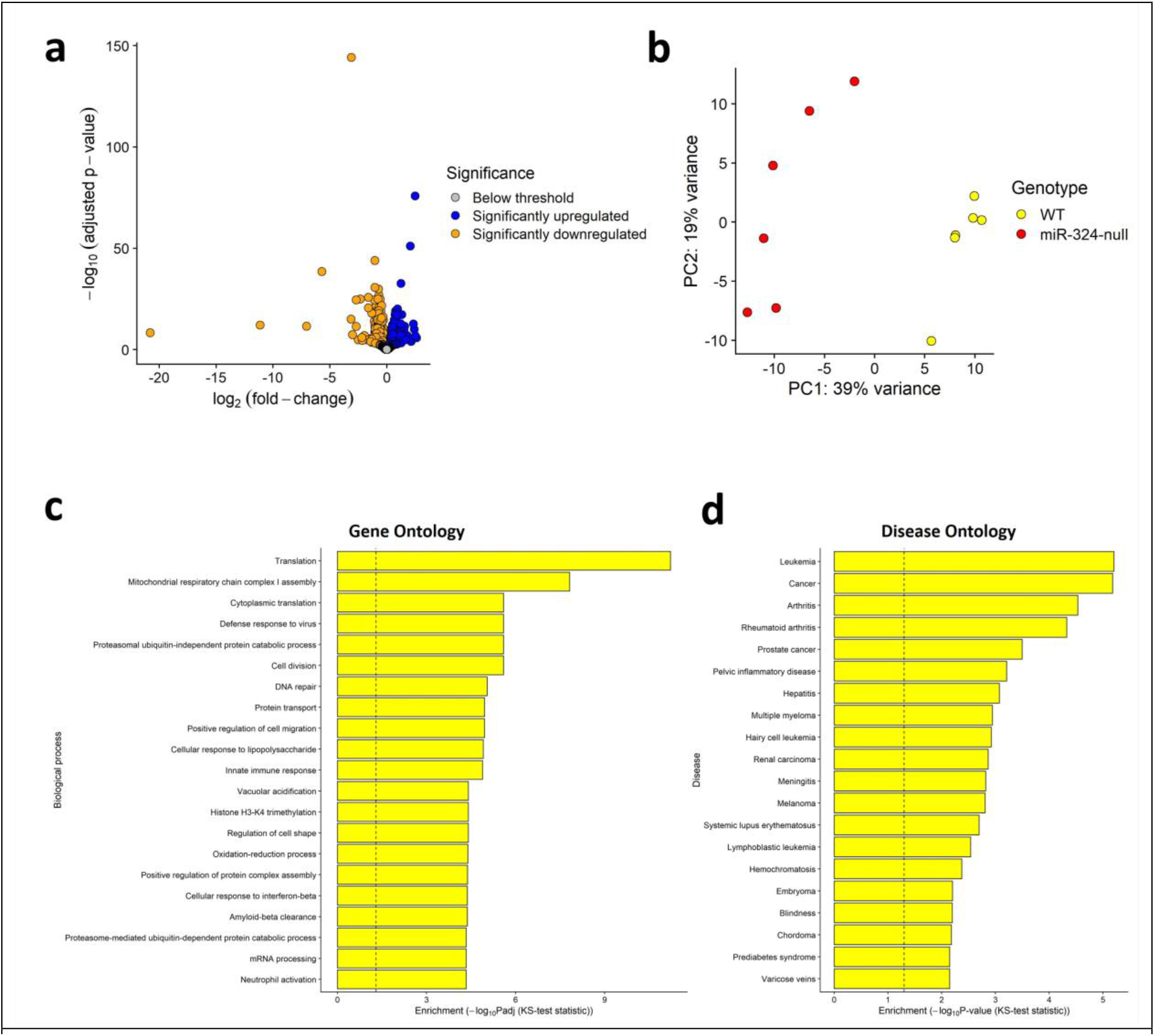
The miR-324-null osteoblast transcriptome is distinct from that of WT osteoblasts. Osteoblasts were isolated from the leg bones of 6 miR-324-null and 6 WT mice aged 20-weeks and grown to confluency before being cultured for 18 days in osteogenic media. Cells were subsequently lysed and the RNA extracted and sequenced. (**a**) Between miR-324-null and WT osteoblasts, 3505 genes were significantly differentially expressed (adjusted p-value ≤ 0.05). (**b**) Samples segregated well by genotype upon PCA. (**c** and **d**) Terms enriched in miR-324-null osteoblasts using GO and DO enrichment analysis. For each plot, a dotted line indicates the significance threshold used to identify statically significantly enriched terms (adjusted p-value ≤ 0.05, adjusted using a Benjamini-Hochberg multiple-testing correction), utilising a Kolmogorov-Smirnov (KS) test.

Together these results demonstrate that the transcriptome of miR-324-null osteoblasts is distinct from that of WT osteoblasts.

### *Runx2* is a direct target of miR-324-5p in osteoblasts

To elucidate the pathways resulting from lack of miR-324 in osteoblasts, the 1,596 significantly upregulated genes were filtered to include those known to have an impact on bone- or metabolism-related disorders using Disease Ontology (26) (Supplementary Table S1). These were subsequently filtered using the TargetScan miRNA target prediction algorithm and the top targets selected for validation using 3’UTR-luciferase assays (Figure 6a). Four osteoblast putative miR-324 target genes; *Klf7*, *Cxcl12* (which encodes SDF-1), *Top1* and *Runx2* were selected (Figure 6b). Multiple *Cxcl12*

**Figure 6.**
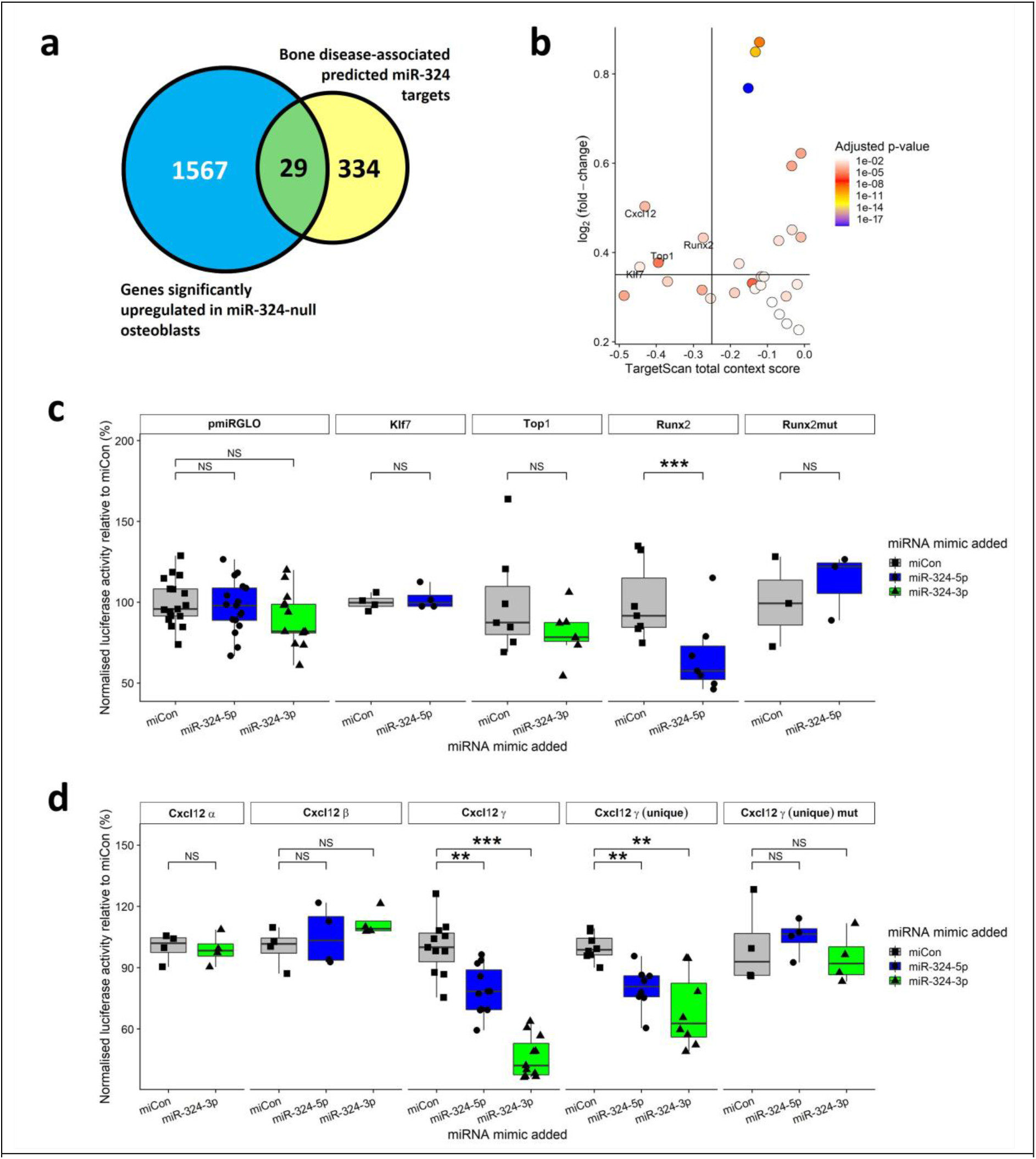
*Runx2* and *Cxcl12* are miR-324 osteoblast targets. (**a**) Of the 1596 genes significantly upregulated between miR-324-null and WT osteoblasts, 29 genes were annotated to bone- or metabolism-related Disease Ontology (26) terms. (**b**) In order to only test the most promising target candidates, thresholds for the TargetScan total context score and the log2 fold-change of < -0.25 and > 0.35, respectively, were utilised (1, 29). Four genes passed these thresholds: *Cxcl12*, *Runx2*, *Top1* and *Klf7*. (**c**) C3H10T1/2 murine cells were transfected with 3’UTR-pmiRGLO constructs for *Runx2*, *Top1* and *Klf7* and either a negative control miRNA mimic (miCon) or a mimic of miR-324-5p or miR-324-3p. Luciferase activity was measured in cell lysates 24 hours post transfection and normalised to control Renilla activity. Values were plotted as a percentage of mean miCon luciferase activity for each construct. The *Runx2* 3’UTR resulted in a significant downregulation of luciferase activity when miR-324-5p mimic was added, and this effect was ablated upon mutation of 2 nt in the predicted miR-324-5p binding site. (**d**) Only the *Cxcl12*-γ and *Cxcl12*-γ (unique), which excludes the part of the *Cxcl12*-γ 3’UTR shared with *Cxcl12*-β, 3’UTRs resulted in a significant down-regulation of luciferase activity when miR-324 mimics were added. When 2 nt in the each *Cxcl12*-γ (unique) predicted binding site of miR-324 were mutated, the ability of the miRNA to repress the luciferase gene upstream was ablated. The means of at least 4 independent experiments were used to test statistical significance, which were calculated from ≥ 3 technical replicates in each independent experiment. A negative control plasmid (pmiRGLO) and negative control mimic (miCon) in addition to a previously validated miR-324 target 3’UTR (as a positive control) were included for each independent experiment. For luciferase assays, paired t-tests were used to assess statistical significance. ***, ** and * represent p-values ≤ 0.001, 0.01 and 0.05, respectively, calculated using Student’s two-tailed paired t-tests.

3’UTR-luciferase reporters were generated since the murine gene can be one of 3 discrete isoforms, *Cxcl12*-α, *Cxcl12*-β and *Cxcl12*-γ, none of which share an identical 3’UTR or all of the same miR-324- 5p or -3p predicted binding sites (Supplementary Figure S4) and all were upregulated in the miR-324- null osteoblasts. *Cxcl12*-α is generally considered to be the canonical isoform (27). Transfection of C3H10T1/2 cells with the reporters and miR-324 mimic or negative control revealed that *Klf7, Top1*, *Cxcl12*-α and *Cxcl12*-β were not direct targets of miR-324. However, the *Runx2* and *Cxcl12*-γ 3’UTRs displayed significant repression of the upstream luciferase gene when miR-324-5p (for *Runx2*), or miR-324-5p and -3p (for *Cxcl12*-γ) were included (Figures 6c and 6d). Since the other *Cxcl12* 3’UTR- luciferase reporters were not repressed by miR-324 we hypothesised that miR-324-5p and –3p were targeting *Cxcl12*-γ at sites unique to this 3’UTR, although this sequence was not predicted to contain a miR-324-3p binding site by TargetScan. However, the miRNA target prediction algorithm, miRmap (28), which allows for limited G-U base pairing between the predicted binding site and the miRNA seed, identified a miR-324-3p binding site within the 3’UTR unique to *Cxcl12*-γ. Indeed a 3’UTR- luficerase reporter of just the unique *Cxcl12*-γ 3’UTR sequence was repressed by both miR-324-5p and miR-324-3p. The miR-324-mediated repression of both *Runx2* and *Cxcl12*-γ (unique) were abolished when 2 nucleotides in each miR-324 binding site (of which *Cxcl12*-γ contained three) were mutated (Figures 6c and 6d).

### miR-324-5p and *Runx2* regulate the balance between osteogenesis and adipogenesis

Both *in vivo* and *ex vivo*, miR-324-null samples showed increased bone formation and *in vivo* almost complete lack of lipid droplets within the tibial bone marrow (Figures 2-4). Considering these results and that we identified *Runx2*, an essential osteogenic transcription factor, as a direct miR-324-5p target (Figure 6), we hypothesised that the miR-324-null high bone mass and reduced lipid phenotypes were driven directly by *Runx2* dysregulation. To investigate the mechanism of altered osteogenesis and adipogenesis, we transfected the immortalised murine MSC-like cell line C3H10T1/2 with miR-324-5p miRNA mimic or hairpin (hp-) inhibitor (as this is the miR-324 arm shown to regulate *Runx2*) and stimulated osteogenesis and adipogenesis using co-differentiation (AdiOst) medium (30). After 14 days of stimulation parallel cell cultures were stained with alizarin red-S or oil red-O staining, for mineralisation and fat deposits, respectively, and the ratio of staining intensity determined. Inhibition of miR-324-5p significantly increased osteogenesis whilst inhibiting adipogenesis and the converse was true with the miR-324-5p mimic (Figure 7a). Mechanistically, inhibition of the miRNA increased *Runx2* but decreased *Pparg* (a master regulator of adipogenesis) expression while the mimic caused an increase in the expression of the latter (Figure 7b). Indeed, overexpression of Runx2 alone (Figure 7c) replicated these results, increasing osteogenesis at the expense of adipogenesis with samples showing a decrease in *Pparg* expression (Figures 7d and e). Overall, these results phenocopy the *in vivo* data and imply that much of the action of miR-324 on osteogenesis occurs through the repressive effect it exerts on Runx2.

**Figure 7.**
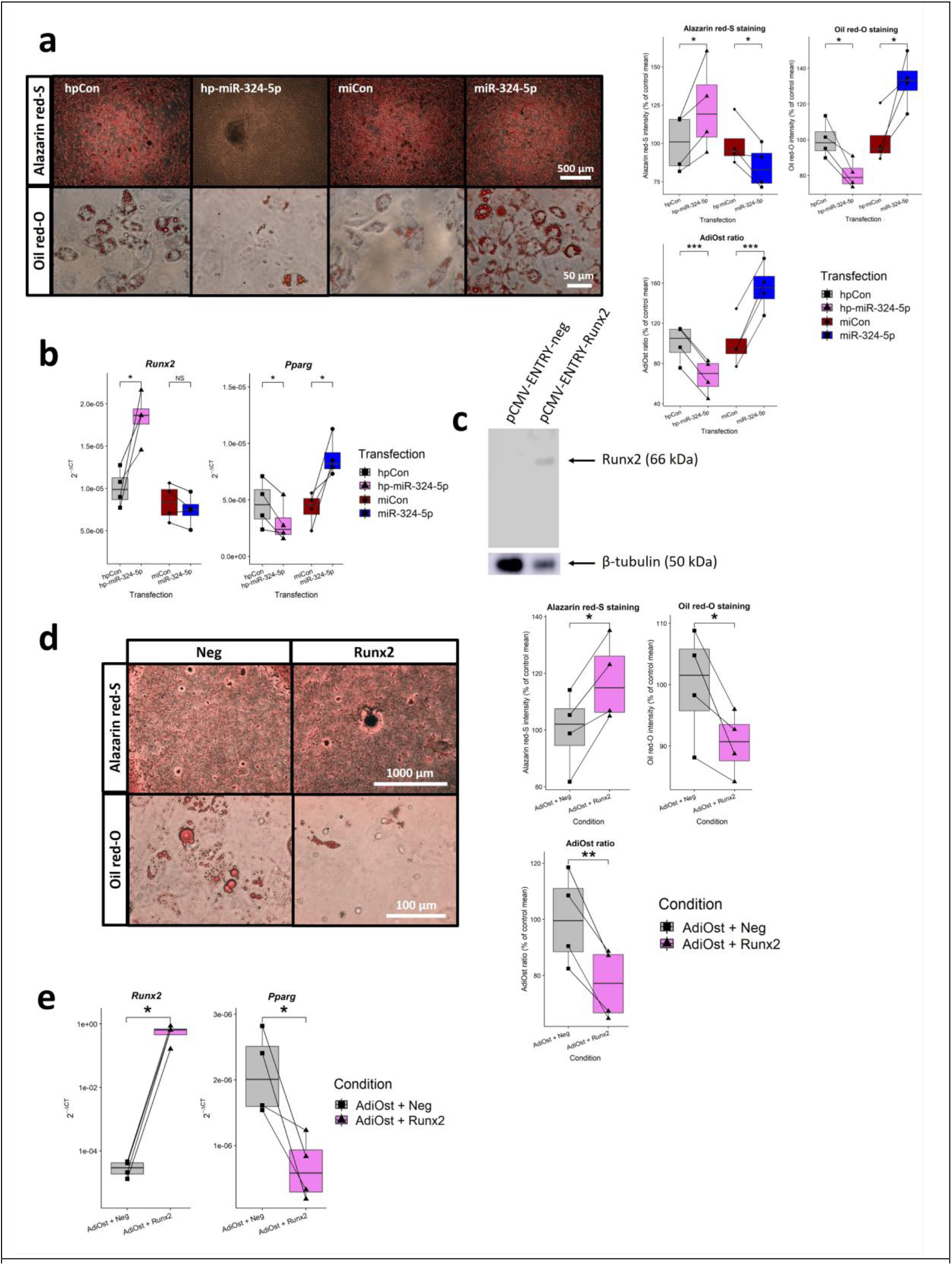
Alteration of miR-324-5p and Runx2 levels modulates the lineage commitment between adipogenesis and osteogenesis in murine C3H10T1/2 cells. C3H10T1/2 cells were transfected either with the negative control miRNA mimic (miCon), miR-324-5p mimic, the negative control miRNA hairpin inhibitor (hpCon) or miR-324-5p hairpin inhibitor (hp-miR-324-5p; at a final concentration of 50 nM). After 18 days of culture in AdiOst media, cells were fixed and stained with alizarin red-S and oil red-O, as indicators of the levels of osteogenesis and adipogenesis, respectively. (**a**) Representative images of C3H10T1/2 cells plated in AdiOst co-differentiation media, transfected with either hpCon, hp-miR-324-5p, miCon or miR-324-5p, and stained either with alizarin red-S or oil red-O. The ratio of oil red-O to alizarin red-S intensity was termed the apidogenesis-osteogenesis (AdiOst) ratio. (**b**) RNA was extracted from each condition and RT-qPCR revealed that inhibition of miR-324-5p significantly increased *Runx2* expression and decreased *Pparg* expression. Overexpression of miR-324-5p significantly upregulated *Pparg* but did not significantly (NS) downregulate *Runx2* expression (p-value = 0.193). (**c**) The miR-324-5p target gene *Runx2* was overexpressed in C3H10T1/2 cells. Western blotting demonstrated that FLAG-tagged Runx2 (detected using an anti-FLAG primary antibody) was overexpressed in transfected cells but not in cells transfected with a negative control plasmid. β-tubulin was used as a protein level normaliser. After Runx2 overexpression and 18 days of co-differentiation the AdiOst ratio was determined (as in **a**) to test whether *Runx2* dysregulation was the mechanism which resulted in increased bone formation in miR-324-null osteoblasts. (**d**) Representative images of cells stained with alizarin red-S (5X magnification) and oil red-O (40X magnification) following 18 days of AdiOst treatment. Cells overexpressing *Runx2* displayed a significant increase in alizarin red-S staining intensity, a significant decrease in oil red-O staining intensity and a significant decrease in the AdiOst ratio. (**e**) RT-qPCR was utilised to demonstrate that overexpression of *Runx2* significantly downregulated *Pparg*. Here, ***, ** and * represent p- values ≤ 0.001, 0.01 and 0.05, respectively, calculated using Student’s two-tailed paired *t*-tests. The means of 4 independent experiments were used to test statistical significance, with 3 technical replicates in each independent experiment. All RT-qPCR results were normalised using the housekeeping gene *18S*.

### The osteoclast and bone marrow macrophage transcriptomes are severely modulated in miR-324-null mice

*In vivo*, osteoblast-mediated bone formation is balanced by osteoclast-mediated bone resorption in healthy individuals. Therefore, the abundance of osteoclasts was also investigated, as the activity of these cells may also be regulated by miR-324. 7- and 14-month miR-324-null tibiae were stained for Tartrate-resistant acid phosphatase (TRAP), a key osteoclast marker, revealing that the mean number of osteoclasts per bone volume was statistically significantly reduced in miR-324-null mice (Figure 8a). The osteoclast surface area per bone surface area was also reduced, although not significantly. These results in tandem with the increased bone formation suggest that the high bone mass phenotype in miR-324-null mice may be due to both reduced bone resorption and increased bone formation. To investigate whether this effect could be replicated *ex vivo*, hematopoietic stromal cells (HSCs) were isolated from miR-324-null and WT bone marrow and stimulated with M-CSF to produce bone marrow macrophages (BMMs). These were subsequently stimulated with M-CSF and RANK-L to induce osteoclastogenesis. TRAP staining of the osteoclast cultures revealed that the lack of miR-324 resulted in fewer osteoclasts, a reduced osteoclast area and fewer nuclei per osteoclast (Figure 8b).

**Figure 8.**
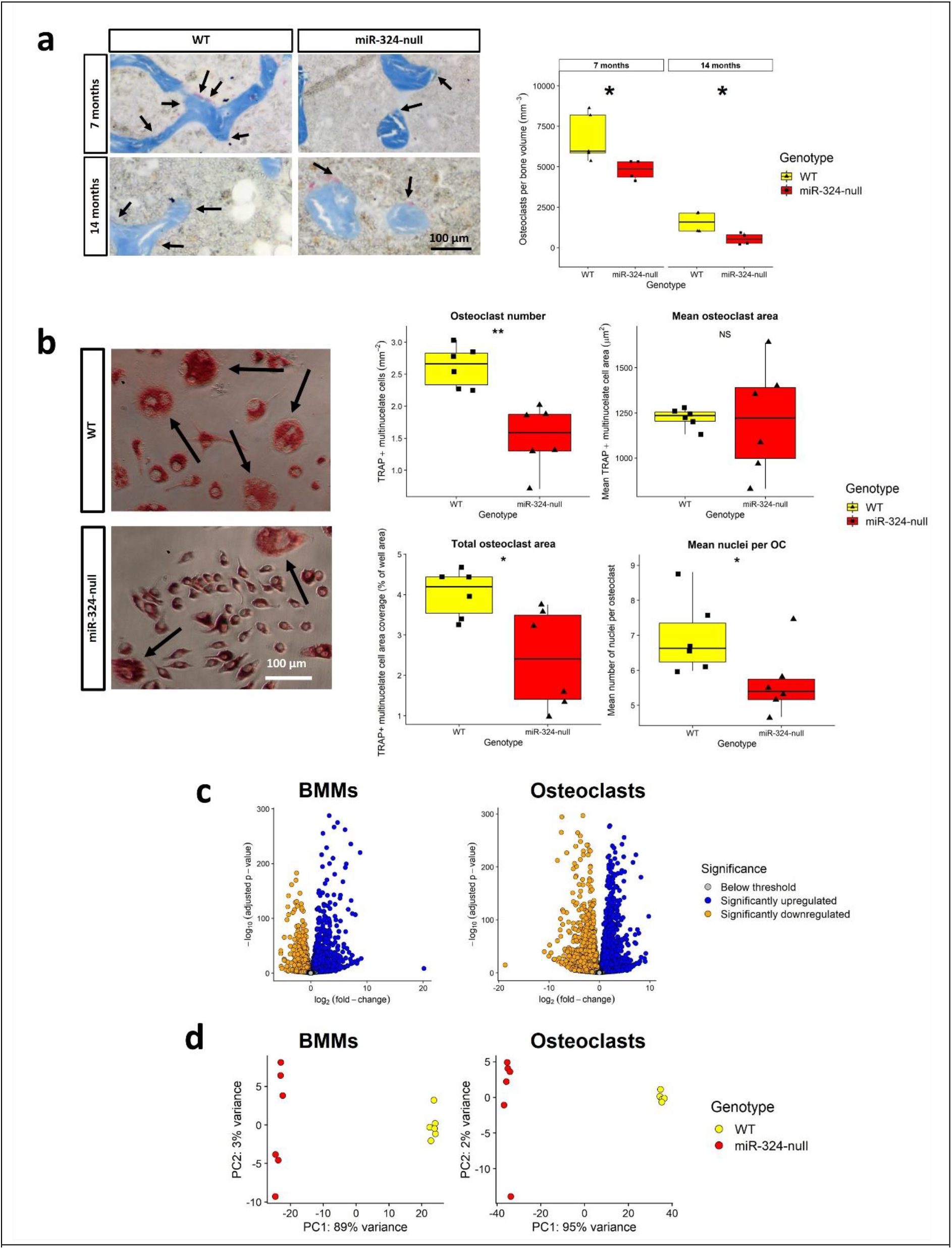
Osteoclastogenesis and the haematopoietic lineage is impaired in miR-324-null mice. (**a**) Formalin-fixed tibial bone sections from miR-324-null and WT mice aged 7- or 14-months were stained for TRAP and analysed using TrapHisto (25). Arrows indicate the osteoclasts. The number of osteoclasts was significantly reduced in miR-324-null samples at both ages. (**b**) HSCs were extracted from the bone marrow of miR-324-null and WT mice and stimulated with 100 ng/ml M- CSF for 3 days, then subsequently with 30 ng/ml M-CSF and 50 ng/ml RANK-L for a further 5 days. miR-324-null samples showed a severe reduction in the number of mature osteoclasts and the mean number of nuclei per osteoclast relative to WT controls. No reduction in mean osteoclast area was observed, although a significant decrease in total osteoclast area was identified in miR-324-null samples. Black arrows on the representative images indicate examples of mature osteoclasts in each genotype. For panels **a** and **b**, statistical significance was assessed using Student’s two-tailed t-tests, where ** and * represent p-values ≤ 0.01 and 0.05, respectively. (c) RNA sequencing of miR-324-null and WT BMMs (M-CSF stimulated) and osteoclasts (M-CSF and RANK-L stimulated) gave 7021 and 8865 differentially expressed genes, respectively, (adjusted p-value ≤ 0.05). Blue and orange points represent significantly upregulated and down-regulated genes, respectively. (d) Both BMM and osteoclast samples segregated by genotype by PCA.

To identify the cause of this osteoclastogenic inhibition, RNA-seq was undertaken in the miR-324- null and WT HSC-differentiate osteoclasts. At a transcriptomic level, miR-324-null osteoclasts were highly distinct from WT osteoclasts; remarkably a total of 8865 genes were differentially expressed between genotypes (43% of all genes detected). This implies that the lack of osteoclastogenesis in miR-324-null samples is probably due to a defect in BMMs or HSCs prior to osteoclast differentiation. In light of this we conducted RNA-seq of miR-324-null and WT BMMs (HSCs stimulated with only M-CSF), in which a total of 7021 genes were significantly differentially expressed in miR-324-null BMMs relative to WT controls (Figure 8c and Supplemental data table 3). Even given these differences the most highly defined cell type using the transcriptomic data from both miR-324-null and WT BMMs was “Macrophages”. In contrast, in the WT osteoclast transcriptomic data “Osteoclasts” was the most highly defined cell type, whereas in miR-324-null osteoclasts this remained as “Macrophages” (Supplementary Table S2). In both experiments, samples segregated by genotype on PCA (Figure 8d). Of note, included in the significantly down- regulated genes in both data sets were *Nfatc1* and *Tnfrsf11a*, the genes encoding the key osteoclastogenesis transcription factor and the RANK receptor, respectively (31, 32). A number of pathways relevant to osteoclastogenesis were also enriched within the datasets (Supplementary Figure S5a). Interestingly, *Acp5*, which encodes TRAP, and *Mmp9* were downregulated in miR-324- null osteoclasts yet upregulated in the BMMs (Supplementary Figure S5b). The majority of genes dysregulated in both cell types were dysregulated in the same direction in both BMMs and osteoclasts (Supplementary Figure S6).

We next attempted to identify targets in the BMMs, since there were fewer significantly differentially expressed genes. In all, 3698 genes were significantly upregulated (adjusted p-value ≤ 0.05 and log2FC > 0). Using the same target prediction pipeline as for the osteoblast dataset, 42 bone or metabolism disease-associated putative miR-324 targets were significantly upregulated in the BMM dataset (Figure 9a), of which 9 passed the thresholds for the TargetScan total context score and the log2 fold-change (< -0.25 and > 0.35, respectively); *Ccne1*, *Pdgfa*, *Klf7*, *Icam1*, *Bcl2*, *Pin1*, *Ing1*, *Ptgs1* and *App* (Figure 9b). 3’UTR luciferase reporter analysis of each of these genes revealed that *Pdgfa*, *Pin1*, *Ptgs1* and *App* were direct miR-324 targets (Figure 9c), with mutated miR-324 seed-sequences relieving the miRNA mediated repression (Figure 9d). Therefore *Pdgfa*, *Ptgs1*, *App* and *Pin1* are direct miR-324 target genes. Notably, 3 of these 4 miR-324 target genes have been implicated in the process of osteoclastogenesis (33–36). It is therefore likely that the repressive effect miR-324 knockout exerts upon osteoclast differentiation is at least partially due to dysregulation of these 3 genes.

**Figure 9.**
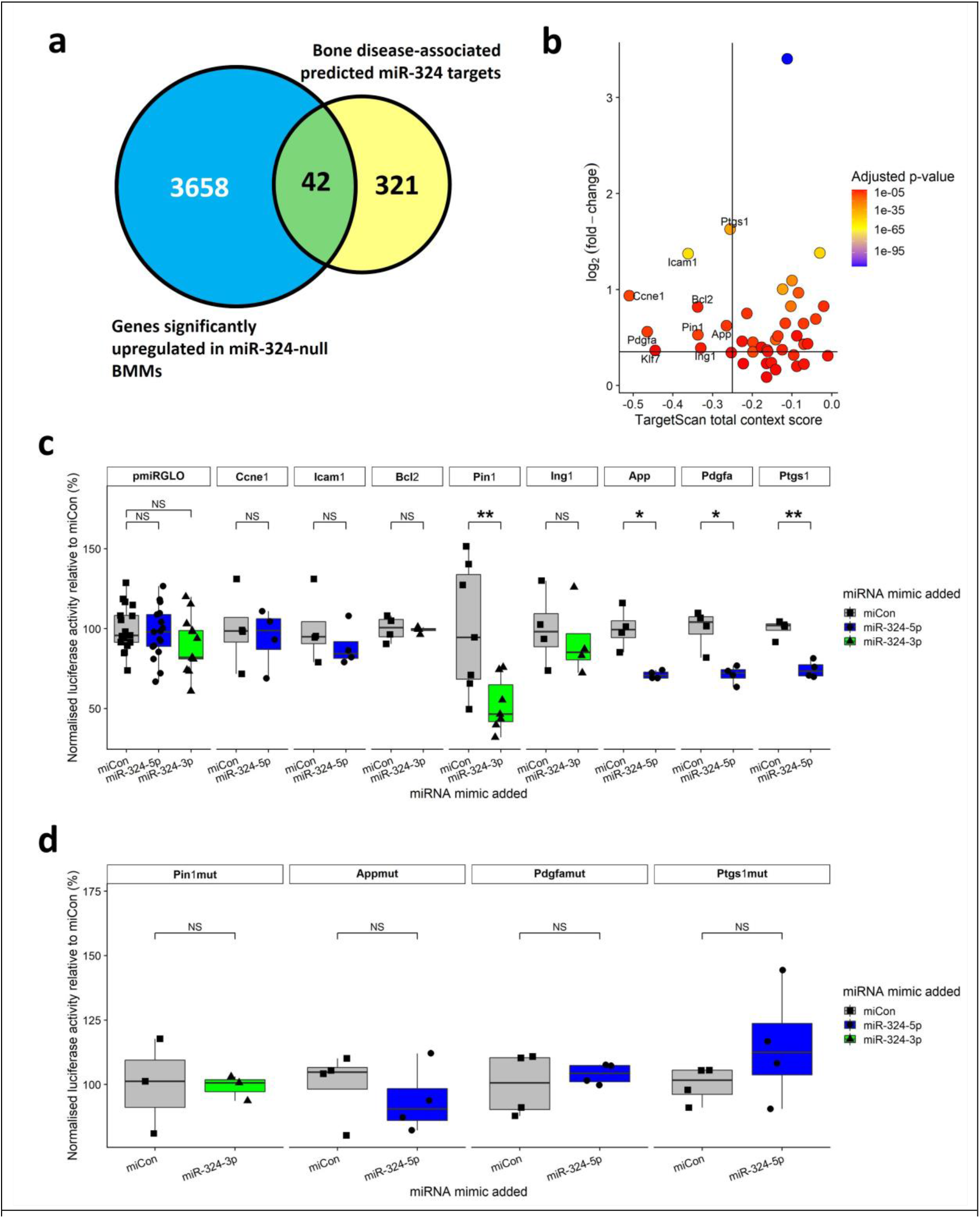
Important osteoclastogenesis genes are direct targets of miR-324. (**a**) Of the 3698 genes significantly upregulated between miR-324-null and WT BMMs, 42 were annotated to bone- or metabolism-related Disease Ontology terms (26). (**b**) 9 genes passed the thresholds of TargetScan total context score and the log2 fold-change of < -0.25 and > 0.35, respectively; *Ccne1*, *Pdgfa*, *Klf7*, *Icam1*, *Bcl2*, *Pin1*, *Ing1*, *Ptgs1* and *App*. (**c**) C3H10T1/2 murine cells were transfected with 3’UTR-pmiRGLO constructs for each putative target gene and either a negative control miRNA mimic (miCon) or a mimic of miR-324-5p or miR-324-3p. After 24 hours Luciferase activity, relative to Renilla, was measured. Values are plotted as a percentage of mean miCon luciferase activity for each construct. The *Pin1, App, Pdgfa* and *Ptgs1* 3’UTRs were found to be targets of miR-324, which were confirmed to be due to direct miR-324 binding through 2 nt mutation of the predicted miRNA-binding site (**d**). The means of at least 4 independent experiments were used to test statistical significance, which were calculated from ≥ 3 technical replicates in each independent experiment. A negative control plasmid (pmiRGLO) and negative control mimic (miCon) in addition to a previously validated miR-324 target 3’UTR (as a positive control – data not shown) were included for each independent experiment. For luciferase assays, paired t-tests were used to assess statistical significance. ***, ** and * represent p-values ≤ 0.001, 0.01 and 0.05, respectively, calculated using Student’s two-tailed paired t-tests.

## Discussion

We initially investigated miR-324 because we identified the miRNA as dysregulated in a miRNA expression screen of normal and osteoarthritic cartilage (15). Having developed the only miR-324- null mouse (20), which develops macroscopically normally, our initial experiments pursued the effects of miR-324 loss on OA development in terms of cartilage destruction, in both surgically-induced and age-onset models. Indeed, OA was worse in the miR-324-null mice. However, chondrocyte transcriptome analysis, both of developing (growth plate) and adult articular cartilage, identified few genes, and very few miR-324 predicted target genes, increased in expression in the miR-324-null cells/tissue. This led us to speculate whether the cause of the increased OA in these mice was due to changes in other joint tissues, especially bone since the miRNA was more abundantly expressed in wild-type bone than cartilage (Supplementary Figure S1). Bone alterations almost certainly play an important role in OA development (37) and the perturbation of genes specifically in bone can alter the susceptibility of mice to OA, as determined by histological examination of cartilage (38). From the bone analysis we found that mice lacking the *Mir324* locus had an increase in trabecular and cortical thickness, with evidence for reduced osteoclastogenesis and increased bone formation.

Numerous studies have described the importance of miRNAs in bone formation and homeostasis (39). The conditional deletion of miRNA species by removal of the miRNA processing enzyme Dicer in osteoblasts (with an Osteocalcin-driven Cre) and osteoclasts (via a CD11b or Cathepsin K-Cre) both result in mice with increased bone mass (40–42), however this phenotype is somewhat timing and Cre-driver dependent (39). Whilst it is not uncommon for a miRNA to affect a component of skeletal biology (39, 43, 44), it is unusual for more than one component of bone remodelling to be altered. For example, miR-185-null mice display increased osteoblast-mediated bone formation (10). Similarly, osteoblast-selective deletion of miR-34b and miR-34c causes increased bone mass (45). However, some miRNAs may affect more than one cell-type in bone. Osteoclast-specific deletion of miR-182 increases post-natal trabecular bone mass (46), while (in zebrafish) the miRNA suppresses osteoblast differentiation (47). miR-21 enhances both osteogenesis and osteoclastogenesis *in vitro* (48, 49), however, although miR-21-null mice display increased bone volume, bone mineral density and trabecular thickness, this appears to be due to dysregulated osteoclast activity as opposed to osteoblast-mediated bone formation. Therefore uniquely, the loss of miR-324 consistently affects osteoblasts and osteoclasts when examined *in vivo*, *in vitro* or *ex vivo*.

Perhaps contrary to our murine observations, a positive correlation has been identified between human miR-324-3p serum abundance and BMD, with miR-324 contributing a group of 8 miRNAs that can be used as a discriminator between controls and patients with low-traumatic fractures [18]. However, these measurements were obtained from human serum and therefore the cellular origin of these serum miRNAs are unclear. A possible explanation for the discordance with our data is that miR-324 is secreted from cells because of its ability to repress bone formation, although this hypothesis requires further investigation to determine conclusively.

miR-324-null mice present with an osteoclast deficiency *in vivo* and impaired osteoclastogenesis *ex vivo*. Over 40% of all genes tested were significantly dysregulated in ex vivo miR-324-null osteoclasts (adjusted p-value ≤ 0.05) and more than half of these were also dysregulated even in BMMs, suggesting the defect is present earlier in the haematopoietic lineage, possibly the HSCs. Conditional deletion of *Dicer* in HSCs confirms that miRNAs are essential for HSC differentiation and downstream activity (50). Specific miRNAs important for haematopoiesis include miR-125a whose overexpression results in a downregulation of HSC apoptosis *in vivo* (50). Meanwhile, ectopic expression of either miR-29a or miR-125b increases HSC proliferation rate and eventually leads to the development of acute myeloid leukaemia (51, 52). The evaluation of miR-324 loss in haematopoiesis would therefore be of interest. It is possible the consequence of miR-324 deletion on haematopoiesis is indirect.

Adipocyte-derived lipids found in bone marrow have been demonstrated to have a supportive role in HSC proliferation (53), reportedly via adipocyte-secreted factors such as adiponectin and leptin (54–56). The observed decreased *in vitro* adipogenesis and the reduced *in vivo* bone marrow lipid accumulation following miR-324 deletion, potentially due to the upregulation of the miR-324-5p target gene *Runx2* (Figure 8), may therefore consequently alter the bone marrow niche leading to haematopoietic dysregulation.

An interesting parallel can be observed between the miR-324-null phenotype and that of sclerostin (*Sost*)-null mice (57). In both, bone formation is increased although this increase is notably greater in the *Sost*-null mice than in miR-324-null mice; trabecular thickness is increased in 5-month-old male miR-324-null mice by 12%, whereas male 5-6.5-month-old *Sost*-null mice display a 146% increase.

Sclerostin is secreted by osteocytes and negatively regulates canonical Wnt signalling pathways to suppress osteogenesis. A specific anti-sclerostin antibody, Romosozumab, is an approved treatment of osteoporosis in postmenopausal women at high risk of bone fractures (58). The Gene Ontology term “Wnt signalling” pathway was significantly enriched in the miR-324-null osteoblast transcriptomic data (adjusted p-value = 7.81 x 10^-3^), due to upregulation of genes such as *Sox9* and Wnt6 (adjusted p-values = 8.61 x 10^-5^ and 2.90 x 10^-5^, respectively). Considering the parallels in phenotype and affected pathways between miR-324-null and *Sost*-null mice, in spite of the difference in magnitude, it would be beneficial in future studies to assess whether *Sost* is differentially expressed in miR-324-null osteocytes and therefore contributes to the observed phenotype.

In summary, we showed that miR-324-null mice displayed an increase in bone mass, in addition to a mild increase in cartilage damage. The mice have a decreased number of adipocyte-derived lipid droplets and osteoclasts, while having an increased activity of osteoblasts. Indeed, the increase in osteoblast function could be attributed to up-regulation of *Runx2*, identified here as a novel miR- 324-5p target. These data also suggest miR-324 is an important driver of MSC commitment to the adipocyte lineage. Perhaps most strikingly, deletion of miR-324 causes a dramatic transcriptome alteration in *ex vivo* cultured macrophages and resultant osteoclasts. Further work is required to determine how and when miR-324 functions during haematopoiesis. However, these phenotypes have posited miR-324 as a novel pharmaceutical target which regulates bone diseases through osteoblasts, osteoclasts and, through the modulation of mesenchymal lineage commitment, adipocytes (summarised in Figure 10).

**Figure 10.**
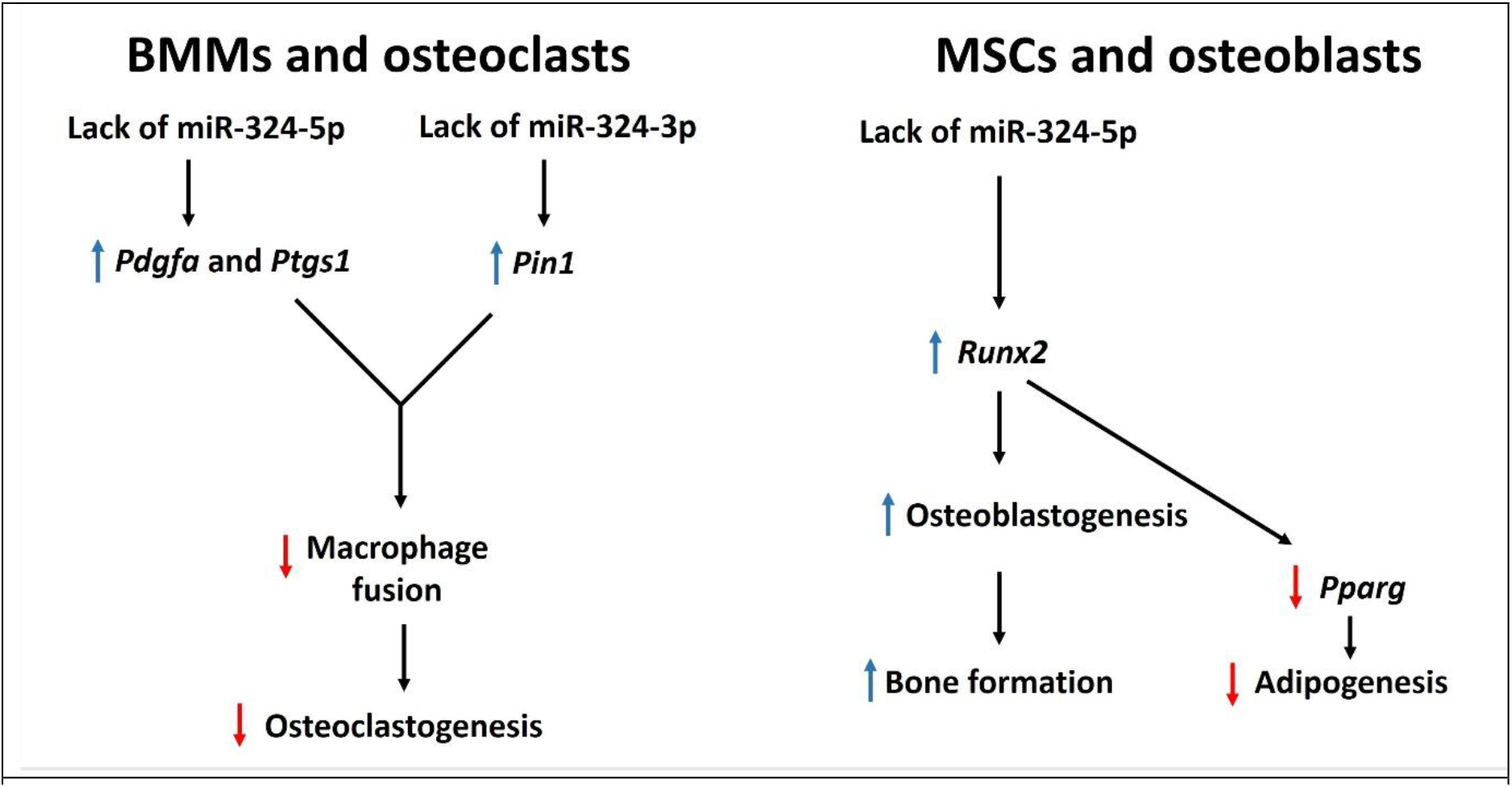
Proposed model of the skeletal system in miR-324-null mice. In the haematopoietic lineage, *Mir324* deletion results in dysregulation of the direct target genes *Pin1* and *Pdgfa*, both of which have been purported to negatively affect osteoclastogenesis (35, 36). In the mesenchymal lineage, dysregulation of the miR-324 target gene *Runx2* results in increased osteogenesis and reduced adipogenesis.

The inhibition of miR-324 offers promise for the treatment of diseases such as osteoporosis, in which bone resorption dominates new bone formation. A deeper understanding of the regulatory role of miR-324 in the musculoskeletal system as a whole is required prior to development of such a treatment, and therefore this should form the basis of future research.

## Methods

### miR-324-null mice

miR-324-null mouse model were generated as previously described (20) and maintained on an inbred C57Bl6 background. All mouse experiments were performed in compliance with the ARRIVE guidelines (https://www.nc3rs.org.uk/arrive-guidelines) (20) under license (P8A8B649A) granted from the Home Office (United Kingdom) in accordance with the guidelines and regulations for the care and use of laboratory animals outlined by the Animals (Scientific Procedures) Act 1986 according to Directive 2010/63/EU of the European Parliament. Protocols were approved by the Animal Ethics Committee of Newcastle University and the Home Office, United Kingdom. Genotyping of the mice was by PCR (Forward primer: 5′-GTGCTGATCTACTCCTCCAACC-3′; reverse primer: 5′- AAATTCACAACTTTGGGGTGAT-3′) of genomic DNA from ear-notches as previously described (20).

Mice were euthanised by cervical dislocation.

### Destabilisation of the medial meniscus (DMM) model

The DMM surgical OA model was performed on the WT and miR-324-null mice essentially as described previously (21). All surgery was performed on male mice at 16-weeks of age. These had their left knee medial meniscus destabilised by transecting the MMTL with a needle blade. The surgical wound was closed with Reflex 7mm wound clips that were subsequently removed seven days post-surgery. Mice were sacrificed 8-weeks post-surgery.

### Histopathological scoring of murine cartilage damage

Left hind limbs of mice were collected and fixed overnight at room temperature (with agitation) in 10% (w/v) neutral-buffered formalin (Sigma-Aldrich). Limbs were washed with distilled water before decalcification with Formical-2000 (American MasterTech Scientific) for 16 hours on a rocker. After decalcification, legs were processed, paraffin-embedded in a patella down orientation, and sectioned and stained with Safranin O/Fast-green as previously described (59). The severity of OA in each section at the medial femoral condyle (MFC) and medial tibial plateau (MTP) of each section was assessed by two independent scorers, blinded to genotype, using the OARSI histopathological scoring system (22). The five highest damage scores given by each scorer at the MFC, the MTP and the five highest scoring sections in terms of summed MFC and MTP scores were averaged, and the means of these averages were used to assess the statistical significance of joint damage between miR-324-null samples and WT controls.

### Extraction of nucleic acids from murine tissues and cells

Total RNA was extracted from murine osteoblasts, osteoclasts and BMMs using the mirVana miRNA Isolation Kit (ThermoFisher Scientific), following the manufacturer’s protocols. Costal cartilage chondrocyte RNA was isolated after initially isolating cells from dissected individual ribs as previously described (60). RNA from medial knee cartilage caps was isolated as described (61). RNA concentration and purity was measured using an Agilent Bioanalyzer 2100 if used for RNA-sequencing. For cells cultured in 96-well plates, the Cells-to-cDNA II Kit (ThermoFisher Scientific) was used for RNA isolation with cDNA synthesis as described (62). Genomic DNA was extracted from murine C3H10T1/2 cells using the PureLink Genomic DNA Mini Kit (ThermoFisher Scientific).

### Real-time reverse transcriptase quantitative PCR (RT-qPCR)

cDNA synthesis was performed using MMLV reverse transcriptase and random hexamers according to the manufacturer’s protocol (ThermoFisher Scientific, Loughborough, UK). For miRNAs, cDNA was synthesised from total RNA using TaqMan® MicroRNA Reverse Transcription (ThermoFisher Scientific) according to the manufacturer’s instructions. TaqMan RT-qPCR was performed, and gene expression levels calculated as previously described (15, 62). Primers/Assay are listed in Supplementary Table S3a, as are the Universal Probe Library probes used (Roche) where applicable. MiRNA expression was normalised to *U6* while other genes of interest (GOI) were normalised to *18S*.

### RNA sequencing

Prior to RNA sequencing (RNA-seq), RNA samples were purified using the DNA*-free*™ DNA Removal Kit (ThermoFisher Scientific) to remove any DNA contamination. Sequencing libraries were prepared using the TruSeq Stranded mRNA sample preparation kit (Illumina) following the manufacturer’s protocol and sequenced with an Illumina NextSeq 500 or NovaSeq. Kallisto (63) was used for pseudo-alignment and quantification, against the mouse GRCh38 (release 92) transcriptome (64).

Mapped transcripts were converted to a gene level using Tximport (65) and DESeq2 (66) was used to calculate p-values and log2 fold-changes (logFC), using the Benjamini-Hochberg method to adjust for multiple testing. EnrichR was utilised to define cells present in miR-324-null and WT osteoclast and BMM samples, using the PanglaoDB Augmented 2021 database with the top 500 most highly expressed genes with human orthologues, determined by average normalised count (67, 68).

### Prediction of novel miR-324 targets

Genes upregulated (adjusted p-value ≤ 0.05 and log2 fold-change > 0.35) in miR-324-null osteoblast and BMM RNA-seq samples were filtered using TargetScan release 7.2 (1) to predict targets of miR- 324-5p and -3p with a total context score of < -0.25. These genes were filtered for those annotated to bone- and metabolism- related DO terms (26) (Supplementary Table S1). The 3’UTRs of genes that passed all filters and thresholds were subsequently tested for direct interaction with miR-324-5p or - 3p using 3’UTR-luciferase assays. miRmap (28) identified an additional miR-324-3p binding site in *Cxcl12*-γ by allowing for a non-canonical G-U base pairing.

### Construction of 3’UTR luciferase reporter plasmids

Fragments of the 3’UTR regions of predicted miR-324 targets were PCR amplified (using Phire Hot Start II DNA Polymerase; ThermoFisher Scientific) from murine genomic DNA or osteoblast cDNA and cloned by In-Fusion HD cloning (Takara Bio) into the pmiRGLO luciferase reporter plasmid (Promega) previously linearised by *Xho*I restriction digestion, essentially as previously described (69).

Oligonucleotides are presented in Supplementary Table S3b. Mutant 3’UTR constructs, where 2 nucleotides within the predicted miR-324 binding sites were mutated, were cloned from gBlocks (Integrated DNA Technologies; Supplementary Table S3c) or for *Ptgs1* generated by site-directed mutagenesis using the QuikChange Lightning Site-Directed Mutagenesis Kit (Agilent Technologies). (Agilent; F primer: AGAAAGAGTCTCACTATGgAaCCATGGCTGGCCTAGAAC, R primer: GTTCTAGGCCAGCCATGGtTcCATAGTGAGACTCTTTCT (mutagenesis sites shown in lower case)). All constructs underwent confirmation by Sanger sequencing.

### 3’UTR-luciferase assays

Murine C3H10T1/2 cells were cultured in DMEM-complete (medium supplemented with 2 mM L-glutamine, 10% foetal bovine serum (FBS), 100 μg/ml streptomycin and 100 IU/ml penicillin) and plasmid transfected essentially as previously described (15). Four hours post plasmid transfection, media was aspirated and the cells were transfected with 50 nM miCon, miR-324-5p or miR-324-3p mimics (Horizon Discovery) using DharmaFECT 1 (Horizon Discovery) transfection reagent. After 24 hours, luciferase and Renilla levels were quantified using a GloMax-Multi Detection System (Promega) and the Dual-Luciferase Reporter Assay System (Promega) (15). Each condition consisted of ≥3 technical replicates and the mean from ≥3 independent experiments were used to calculate statistical significance using Student’s two-tailed paired *t*-tests.

### Protein extraction and immunoblotting

Murine tissue and cells samples were processed using a 1% (v/v) TritonX lysis buffer, as previously described (20). Lysates were resolved by sodium dodecyl sulfate polyacrylamide gel electrophoresis, transferred to polyvinylidene fluoride membranes (Millipore, Watford, UK) and subsequently probed using the antibodies described in Supplementary Table S4, all essentially as previously described (20). Visualisation was undertaken using HRP-conjugated secondary antibodies (Dako; used at 1:1000 dilution) and Immobilon Western Chemiluminescent HRP Substrate (Merck Millipore) and subsequent quantification was performed using Fiji (70).

### Micro-computed tomography (μCT)

After mice were sacrificed by cervical dislocation, their right hind legs were fixed overnight in 10% (w/v) neutral-buffered formalin (Sigma-Aldrich) before transfer to 70% (v/v) ethanol. Limbs were analysed by μCT to examine calcified tissues using a SkyScan 1272 (Bruker, Belgium; 0.5 aluminium filter, 50 kV, 200 μA). Cortical bone was scanned at a voxel size of 9.0 µm, with a 0.5° rotation angle. Trabecular bone was scanned at a voxel size of 4.5 µm using a 0.3° rotation angle. NRecon (Bruker) and DataViewer (Bruker) were used to construct cross-sectional slices or to generate 3D computational models, respectively. CTAn (Bruker) was used to quantify metrics, essentially as previously described (71). For trabecular tibial and femoral measurements, a volume of interest (VOI) encompassing the region 90-990 µm below or above (respectively) the growth plates were used. For measuring tibial cortical bone, a VOI positioned 450-1350 µm above the tibiofibular junction was analysed, whereas for femoral cortical bone, a 900 µm VOI at the vertical midpoint of the femur was analysed. To measure the bowing of the leg bones, the horizontal distance between the fibula and tibia at the midpoint of the tibia (calculated at the exact vertical midpoint between the tibial plateau and the point at which the fibula joins the tibia) was measured on the transaxial plane. Bone tissue mineral density (TMD) was calculated from cortical scans using CtAn (Bruker), calibrated against phantom rods of known calcium hydroxyapatite densities of 0.25 and 0.75 g‧cm^-3^ (Bruker).

### Histology and histomorphometry

Mice underwent intraperitoneal injections (0.1 ml/10 g body weight) of alizarin red-S (3 mg/ml) and then calcein (2 mg/ml) 7 and 1 day prior to sacrifice. After analysis by μCT (above), the tibiae were embedded in methyl methacrylate (MMA) and 5 µm sections cut with a tungsten steel knife on a motorised rotary microtome (Leica). Sections were stained either with Goldner’s Trichrome to analyse osteoid and lipid droplets or counterstained with calcein blue to quantify the amount of bone formed between the injected alazarin red-S and calcein, essentially as previously described (71). Images were captured using an Axioscan Z1 slide scanner (Zeiss) and histomorphometry performed using the OsteoidHisto and CalceinHisto open-source image analysis programs (25). A mean average of 3 sections/mouse was used to represent N = 1.

### Osteoblast and haematopoietic stromal cell isolation and culture

Osteoblasts were isolated from the calvariae of 3-day-old mouse pups essentially as previously described (71). Briefly, dissected calvariae were digested with collagenase type I (Sigma-Aldrich) and the isolated cells cultured in α-MEM (Gibco) supplemented with 2 mM L-glutamine, 10% FBS, and antibiotics. A separate population of osteoblasts and haematopoietic stromal cells were isolated from leg bone chips and marrow respectively, dissected from the long bones of 20-week-old mice. The epiphyses were removed to expose the marrow, which was flushed from the bones by centrifugation and cultured directly with α-MEM (supplemented as above but also with 100 ng/ml M-CSF for 3 days to obtain M-CSF-dependent bone marrow macrophages (BMMs) (71)). The remaining bone was cut into small chips of approximately 2 mm^3^ and collagenase digested for 1 hour. Subsequently, the bone chips were washed with PBS and cultured in supplemented α-MEM and cultured for approximately 6 weeks until semi confluent layers of osteoblasts were present. Osteocytes were also isolated from bone chips, as previously described (72).

### Osteoblast proliferation and alkaline phosphatase assays

For cell proliferation and alkaline phosphatase assays, osteoblasts were cultured in complete DMEM for 24 hours. Proliferation was quantified after 2.5 hours culture with Alamar Blue (ThermoFisher Scientific) following the manufacturers protocol. For alkaline phosphatase assays the cells were formalin-fixed, freeze-thawed and incubated with 20mM paranitrophenyl phosphate (PNPP (ThermoFisher Scientific) which is dephosphorylated by alkaline phosphatase to produced paranitrophenol (PNP), the absorbance of which was measured at 405 nm after 30 minutes. PNP levels were normalised against cell proliferation measurements, all essentially as described (71).

### Osteogenesis assays and alizarin red-S staining in murine osteoblasts

Osteoblasts underwent alizarin red-S staining, adapted from previously described methodology (71). Briefly, cells were cultured in osteogenic medium (α-MEM-complete, supplemented with 2 mM β- glycerophosphate (BGP) and 50 µg/ml L-ascorbic acid (both Sigma-Aldrich)). Calvariae and bone-chip osteoblasts were incubated for 7 days or 18 days, respectively, before being formalin-fixed, stained with 40 mM alizarin red-S for 30 minutes and imaged with an Axiovert 200 Inverted microscope (Zeiss). The alizarin red-S was extracted using 10% (w/v) cetylpyridinium chloride, the absorbance of which was measured 562 nm. Parallel cell cultures were either lysed for protein or RNA using previously described methods.

### miR-324 titration and reintroduction in miR-324-null osteoblasts

miR-324-null and WT calvarial osteoblasts were isolated and cultured as described earlier. miR-324- null osteoblasts were transfected (DharmaFECT 1) with miR-324-5p and -3p mimics at final concentration ranging from 5 nM to 5 pM for 24 hours (Supplementary Figure S3), and subsequently stimulated with osteogenic media for 7 days. Cells transfected with miRIDIAN microRNA Mimic Negative Control #2 (miCon; Horizon Discovery) were included as a negative control and WT osteoblasts as a positive control. RT-qPCR for miR-324-5p and -3p was undertaken as described. Optimal concentration was determined as the level equivalent to the WT osteoblasts and was used to transfect further cultures of miR-null osteoblasts for which gene expression analysis and alizarin red-S staining were undertaken after 7 days of stimulation with osteogenic media (as described above).

### Osteoclastogenesis assays and TRAP staining

M-CSF-dependent BMMs were cultured in α-MEM-complete, supplemented with 50 ng/ml RANK-L and 30 ng/ml M-CSF (Peprotech) to stimulate osteoclastogenesis. Equivalent BMM control cultures were stimulated with just M-CSF. After 5 days, RNA was extracted or cells were fixed for Tartrate-resistant acid phosphatase (TRAP) staining. For TRAP staining, cells were formalin-fixed and assayed using the Acid Phosphatase, Leukocyte (TRAP) Kit (Sigma-Aldrich) following the manufacturer’s instructions. Cells were imaged with an Axiovert 200 inverted microscope (Zeiss) and TRAP-positive cells (with ≥3 nuclei) in each sample were counted in addition to the number of nuclei in each of these cells. Fiji (70) was used to measure the total area covered by TRAP-positive cells with ≥3 nuclei, allowing calculation of the mean osteoclast area. All osteoclastogenesis assays were performed in triplicate and the mean of the triplicates was treated as N = 1 for each mouse from which cells were isolated.

### Construction of negative control overexpression plasmid

A Myc-DDK (FLAG)-tagged coding DNA sequence (CDS) clone of murine *Runx2* was purchased from Origene within the pCMV-ENTRY overexpression vector (catalogue number MR227321). To produce a negative control plasmid containing no CDS (pCMV-ENTRY-neg), the pCMV-ENTRY-*Runx2* plasmid was restriction digested with *Bam*HI and *Xho*I and recircularised by In-Fusion HD cloning a synthesised gBlock (Supplementary Table S3c).

### Adipo-osteogenesis in murine C3H10T1/2 mesenchymal stromal cells

Adipo-osteogenesis (AdiOst) co-differentiation assays were undertaken using murine C3H10T1/2 cells. Cells were transfected with either 50nM microRNA Hairpin Inhibitor Negative Control #1 (hpCon), hp-miR-324-5p hp-miR-324-3p, miCon, miR-324-5p mimic, miR-324-3p mimic (all Horizon Discovery) using DharmaFECT 1, or with the plasmids pCMV-ENTRY-*Runx2* or pCMV-ENTRY-neg using FuGene HD. 24 hours post-transfection cells were stimulated with AdiOst co-differentiation media (α-MEM-complete supplemented with 100 nM dexamethasone, 2 mM BGP and 50 µg/ml L-ascorbic acid (30, 71)) for 18 days. From the experimental wells, either protein or RNA (Cells-to-cDNA II Kit) was extracted or the cells were stained with alizarin red-S (as described earlier) or oil red-O, undertaken as previously described (73); following oil red-O staining, cells were washed, imaged with an Axiovert 200 inverted microscope (Zeiss), and destained using 100% (v/v) isopropanol. The stain intensity was quantified by measuring the absorbance at 500 nm.

### Statistical analysis

Unless indicated, the Shapiro-Wilk test was used to assess normality of data. Statistical significance was assessed using Student’s two-tailed *t*-test (all unpaired unless indicated otherwise in the figure legends) for single comparisons or analysis of variance (ANOVA) for testing the effects of multiple variables on a continuous variable output; the test used for each comparison is indicated in the corresponding figure legend. All data analysis and statistical calculations were performed using R version 3.6.2 (74). Relevant R packages used for statistical analysis are cited.

### Data availability

All RNA-seq data is publicly available at the NCBI Gene Expression Omnibus under the accessions GSE226615, GSE234682 and GSE233809.

## Supporting information

rib chondrocytes

knee chondrocytes

bone cell

## Acknowledgements

For the purpose of open access, the author has applied a Creative Commons Attribution (CC BY) licence to any Author Accepted Manuscript version arising from this submission.

