## supplemental_figures for "miR-324 mediates bone homeostasis through the regulation of osteoblast and osteoclast differentiation and activity"

### Supplementary Data

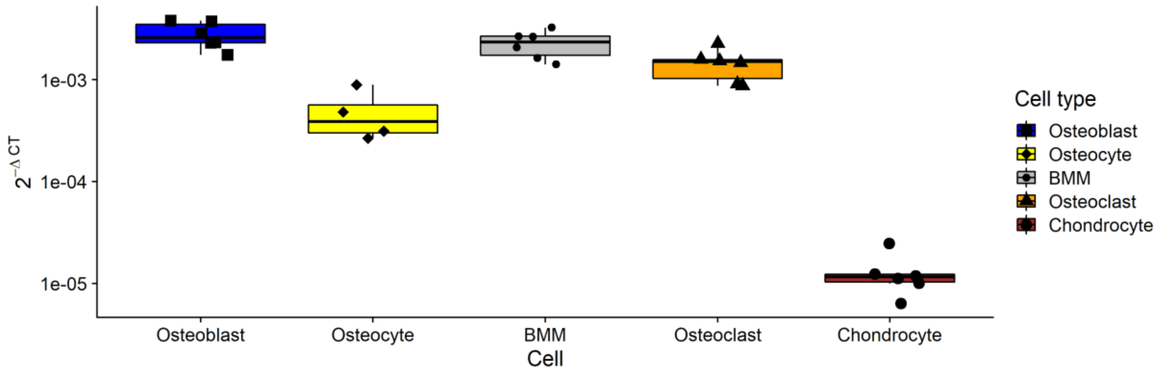


**Supplementary Figure S1 - miR-324 is far more abundant in murine bone cells (bone marrow-derived macrophages (BMMs), osteoblasts, osteoclasts and osteocytes) than in murine cartilage.** BMMs were cultured from WT murine bone marrow stimulated with M-CSF. Osteoclasts were generated from BMMs by further stimulating BMMs with M-CSF in addition to RANK-L. Osteoblasts grown to confluency out of murine leg bone chips (for approximately 6 weeks), before being cultured for 18-days in osteogenic media. Osteocyte RNA was isolated by successive digestions of murine leg bone chips with EDTA and collagenase, followed by grinding the remaining bone chips and extracting total RNA. Chondrocyte RNA was isolated by grinding murine knee cartilage and extracting total RNA. Abundance of miR-324 in each cell type was determined by RT-qPCR and normalised against *U6*. Each point represents the abundance of miR-324 in tissue from a single mouse. For BMMs, osteoclasts and osteoblasts, each point is the mean value from ≥4 technical replicates.

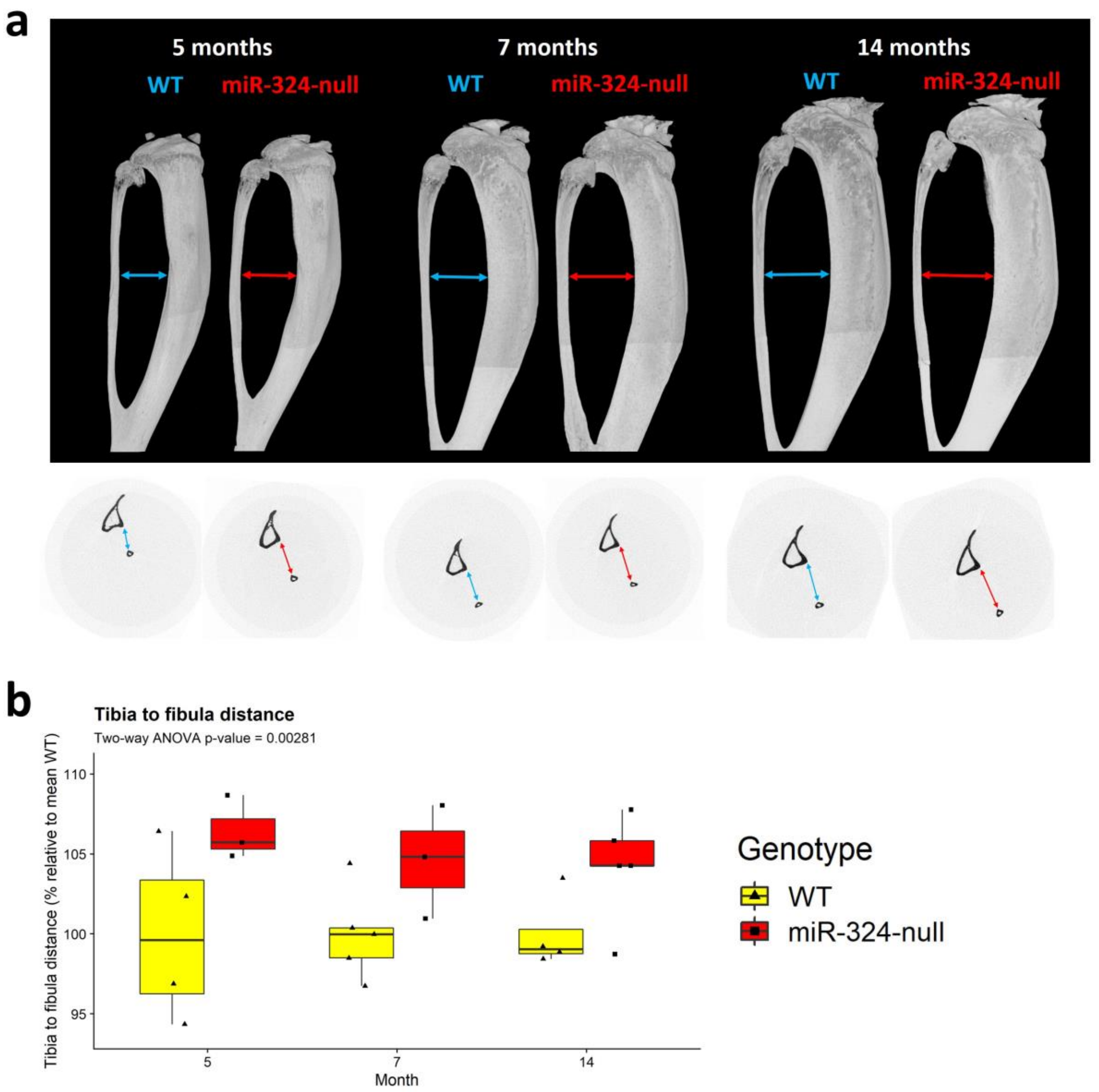


**Supplementary Figure S2 - miR-324-null mice display a tibial bowing phenotype.** (**a**) Representative tibial µCT reconstructions are shown at each time point for miR-324-null or WT mice, illustrating the increased bowing observed in miR-324-null bones. The blue arrow at each age indicates the horizontal distance between the tibial midpoint and the fibula in WT mice, whereas the red arrow indicates the distance in miR-324-null mice, which is increased. Transaxial slices are also shown using the same colours. (**b**) The distance between the tibia and fibula at the tibial midpoint is increased in miR-324-null mice relative to WT controls. Statistical significance overall across time points was assessed using two-way ANOVA and Tukey's HSD post-hoc tests were used to assess statistical significance at individual time points, although no individual time point achieved statistical significance in post-hoc testing.

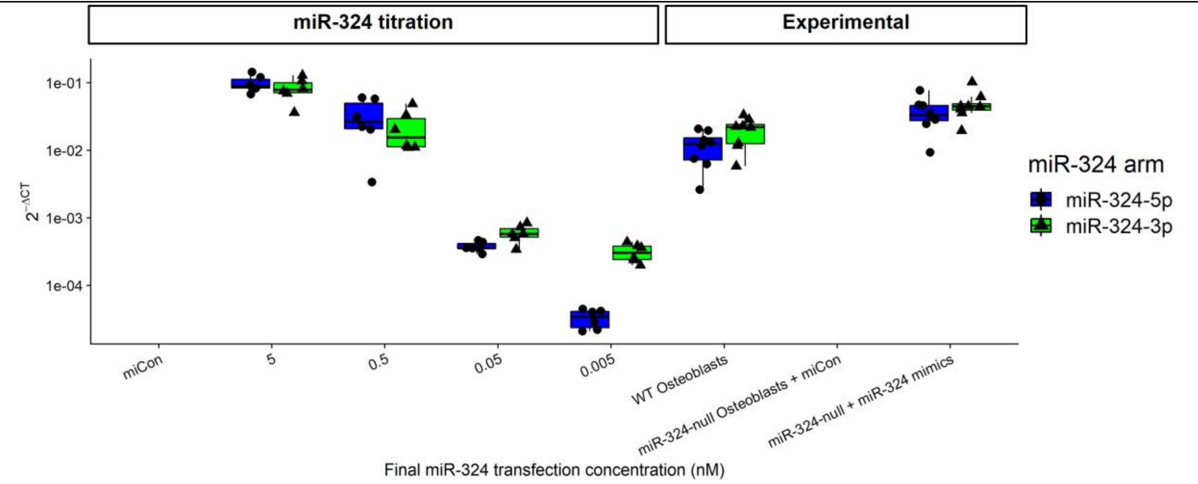


**Supplementary Figure S3 - Titration of the final transfection concentrations of miR-324 -5p and -3p mimics in order to restore each miR-324 arm to physiological osteoblast levels.** miR-324-null osteoblasts were transfected with 50 nM (final concentration) of miCon as a negative control, in addition to 4 test concentrations of each of miR-324-5p and -3p; 5 nM, 0.5 nM, 0.05 nM and 0.005 nM. WT osteoblasts were also included as a positive control. The optimum final transfection concentration to restore miR-324 to near physiological levels was identified as being 0.5 nM for each arm. The transfection of miR-324-null osteoblasts with 0.5 nM miR-324 mimics was subsequently repeated and compared to WT osteoblasts and miR-324-null osteoblasts transfected with 0.5 nM miCon.

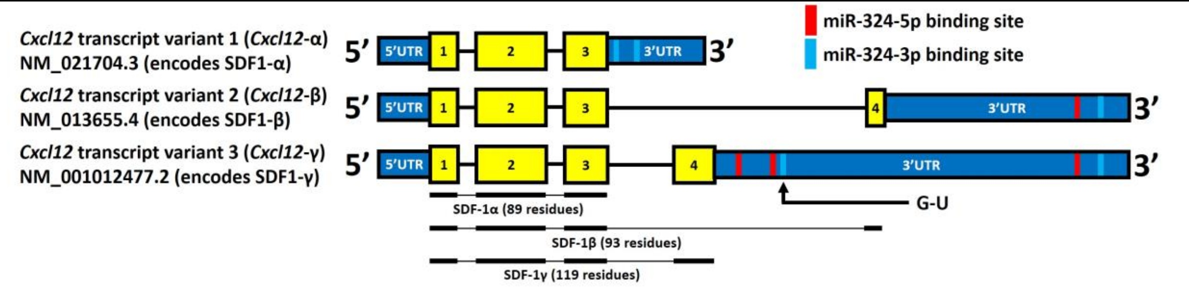


**Supplementary Figure S4 - Isoforms of *Cxcl12*.** There are 3 distinct isoforms of *Cxcl12*, each of which possess distinct 3'UTRs and encode distinct proteins; *Cxcl12*-α, -β and -γ encode SDF-1α, -β and -γ, respectively. All isoforms possess the exons within *Cxcl12*-α, whereas -β and -γ each possess an additional unique exon. For each isoform, the locations of miR-324-5p and -3p binding sites are shown in red and blue, respectively. One of the predicted miR-324-3p binding sites in the *Cxcl12*-γ isoform is predicted to bind miR-324-3p through a non-canonical G-U base pairing (indicated on schematic), and this site is indicated on the diagram.

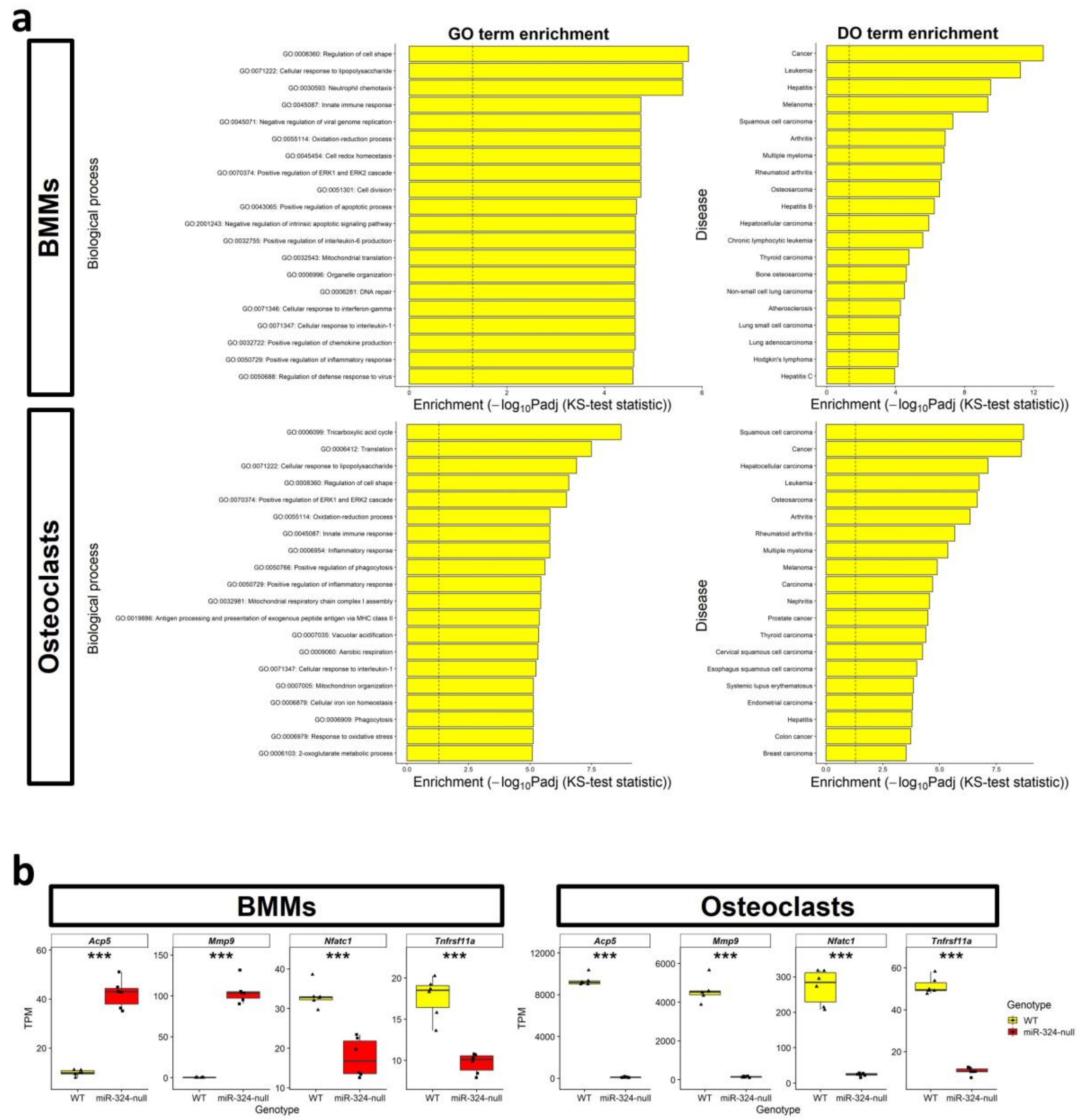


**Supplementary Figure S5 - Important osteoclastogenesis genes are dysregulated in miR-324-null BMMs and osteoclasts.** (**a**) GO and DO enrichment of RNA-seq results revealed that genes annotated to the 'Positive regulation of ERK1 and ERK2 cascade' and genes annotated to 'Arthritis', respectively, were significantly enriched in both the BMM and osteoclast experiments. These have strong links to osteoclast biology in the literature (He et al., 2011; Sato and Takayanagi, 2006). (**b**) Both miR-324-null BMMs and osteoclasts displayed a severe downregulation of the key osteoclastogenesis genes *Nfatc1* and *Tnfrsf11a*. *Tnfrsf11a* is annotated to the "Regulation of ERK1 and ERK2 cascade" GO term and both *Tnfrsf11a* and *Nfatc1* are annotated to the "Arthritis" DO term. *Acp5* and *Mmp9* were also severely downregulated in miR-324-null osteoclasts relative to WT controls, but unexpectedly were significantly upregulated in miR-324-null BMMs. Here, *** indicates adjusted p-values ≤ 0.001, computed as part of RNA-seq analysis.

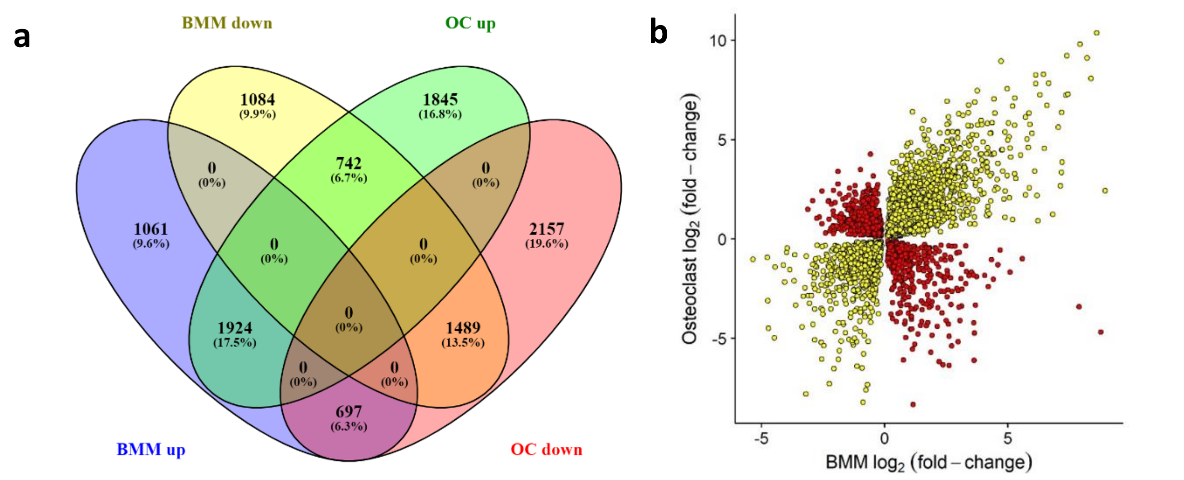


**Supplementary Figure S6**. **Overlap and correlation of the significantly differentially expressed genes in both miR-324-null BMMs and osteoclasts.** More than half of the genes dysregulated in miR-324-null BMMs are also dysregulated in miR-324-null osteoclasts. The total number of genes dysregulated in BMMs totals at 7023, whereas in osteoclasts the number comes to 8882 genes (34.1% and 43.2% of all genes tested, respectively). (**a**) Venn diagram of the overlaps of significantly differentially expressed genes between the BMM and osteoclasts (OCs), segregated into up and down regulated in each cell type. (**b**) A scatter plot comparing the $\log_2$ fold-changes in BMMs and OCs of genes dysregulated in both cell types. Yellow indicates genes with a shared directionality of regulation in both cell types, whereas red indicates genes which are dysregulated in both BMMs and OCs, but not in the same direction. The majority of genes dysregulated in both cell types were dysregulated in the same direction in both BMMs and OCs; 1933 genes (17.5%) were upregulated and 1492 genes (13.5%) downregulated in both cell types. More than 6% of the common significantly differentially expressed genes changed expression in the opposite direction in each case (i.e., BMM up vs OC down and BMM down vs OC up).

| DO term | Total number of genes annotated to term |
|---|---|
| Achondroplasia | 1 |
| Arthritis | 546 |
| Bilirubin metabolic disorder | 1 |
| Bone cancer | 3 |
| Bone giant cell tumor | 13 |
| Bone inflammation disease | 3 |
| Bone lymphoma | 1 |
| Bone marrow cancer | 16 |
| Bone marrow disease | 2 |
| Bone osteosarcoma | 46 |
| Chondrocalcinosis | 2 |
| Chondrosarcoma | 62 |
| Clear cell chondrosarcoma | 1 |
| Disease of metabolism | 8 |
| Extraskeletal myxoid chondrosarcoma | 3 |
| Fibrosarcoma of bone | 3 |
| Hypophosphatasia | 1 |
| Idiopathic juvenile osteoporosis | 2 |
| Inherited metabolic disorder | 1 |
| Iron metabolism disease | 1 |
| Juvenile rheumatoid arthritis | 3 |
| Mesenchymal chondrosarcoma | 2 |
| Metabolic syndrome X | 20 |
| Mitochondrial metabolism disease | 11 |
| Myxoid chondrosarcoma | 3 |
| Osteoarthritis | 183 |
| Osteochondritis dissecans | 15 |
| Osteochondrodysplasia | 5 |
| Osteogenesis imperfecta | 6 |
| Osteomalacia | 7 |
| Osteomyelitis | 15 |
| Osteonecrosis | 26 |
| Osteopetrosis | 13 |
| Osteopoikilosis | 1 |
| Osteoporosis | 90 |
| Osteosarcoma | 214 |
| Osteosclerosis | 6 |
| Paget's disease of bone | 14 |
| Pediatric osteosarcoma | 1 |
| Primary hypertrophic osteoarthropathy | 1 |
| Psoriatic arthritis | 6 |
| Reactive arthritis | 15 |
| Renal osteodystrophy | 2 |
| Rheumatoid arthritis | 512 |

| | |
|---|---|
| Rickets | 8 |

**Supplementary Table S1 - Bone- or metabolism-related Disease Ontology terms utilised to filter putative miR-324 target genes.**

**WT BMMs**

| Cell type | Adjusted p-value |
|---|---|
| Macrophages | 8.97E-30 |
| Osteoclasts | 4.88E-23 |
| Microglia | 7.64E-23 |
| Kupffer Cells | 1.36E-17 |
| Alveolar Macrophages | 6.98E-17 |
| Monocytes | 3.23E-18 |
| Red Pulp Macrophages | 5.25E-15 |
| Osteoclast Precursor Cells | 2.52E-12 |
| Langerhans Cells | 5.89E-13 |
| Myeloid-derived Suppressor Cells | 5.88E-12 |
| Dendritic Cells | 3.98E-14 |
| Eosinophils | 1.82E-07 |

**miR-324-null BMMs**

| Cell type | Adjusted p-value |
|---|---|
| Macrophages | 1.11E-28 |
| Microglia | 8.80E-24 |
| Kupffer Cells | 1.28E-20 |
| Alveolar Macrophages | 5.40E-19 |
| Osteoclasts | 4.94E-20 |
| Red Pulp Macrophages | 3.00E-17 |
| Monocytes | 2.10E-18 |
| Myeloid-derived Suppressor Cells | 4.64E-15 |
| Eosinophils | 1.08E-15 |
| Langerhans Cells | 4.63E-15 |
| Dendritic Cells | 1.06E-17 |
| Osteoclast Precursor Cells | 1.68E-13 |

**WT osteoclasts**

| Cell type | Adjusted p-value |
|---|---|
| Osteoclasts | 1.99E-19 |
| Macrophages | 4.23E-14 |
| Microglia | 5.63E-11 |
| Microfold Cells | 9.25E-10 |
| Monocytes | 6.70E-11 |
| Eosinophils | 9.25E-10 |
| Kupffer Cells | 7.22E-10 |
| Myeloid-derived Suppressor Cells | 4.25E-09 |
| Alveolar Macrophages | 3.08E-09 |
| Langerhans Cells | 3.08E-09 |
| Red Pulp Macrophages | 1.38E-07 |
| Megakaryocytes | 3.45E-08 |
| Osteoclast Precursor Cells | 5.62E-07 |

**miR-324-null osteoclasts**

| Cell type | Adjusted p-value |
|---|---|
| Macrophages | 1.18E-28 |
| Kupffer Cells | 1.14E-22 |
| Osteoclasts | 4.46E-22 |
| Red Pulp Macrophages | 2.30E-19 |
| Alveolar Macrophages | 4.32E-20 |
| Microglia | 7.24E-22 |
| Monocytes | 1.90E-18 |
| Eosinophils | 1.19E-14 |
| Myeloid-derived Suppressor Cells | 5.90E-14 |
| Platelets | 1.03E-16 |
| Osteoclast Precursor Cells | 1.86E-12 |
| Neutrophils | 7.83E-14 |
| Megakaryocytes | 1.67E-13 |

**Supplementary Table S2 - miR-324-null osteoclasts are more transcriptomically similar to macrophages than osteoclasts.** EnrichR was utilised to define cells present in miR-324-null and WT osteoclast and BMM samples, using the PanglaoDB Augmented 2021 database with the top 500 most highly expressed genes with human orthologues, determined by average normalised count (Chen *et al.*, 2013; Franzén *et al.*, 2019). The most relevant cell types to osteoclastogenesis are highlighted.

**a**

| Gene | Forward primer | Reverse primer | UPL probe | Amplicon length (nt) |
|---|---|---|---|---|
| *18S* | (Predesigned TaqMan assay used; catalogue number 4333760T) | | | |
| *Alpl* | CTTCGCTCTCCGAGATGGTG | CCCTCATGATGTCCGTGGTC | 31 | 109 |
| *Col1a1* | CAGCGTAGCCTACATGGACC | CGATGACTGTCTTGCCCCAA | 68 | 167 |
| *Col1a2* | GGACACAGTGGTATGGATGGA | TTGACCTGGAGTTCCATTCTC | 64 | 96 |
| *Mir324-3p* | (Predesigned TaqMan assay used; catalogue number 4427975, assay 002509) | | | |
| *Mir324-5p* | (Predesigned TaqMan assay used; catalogue number 4427975, assay 000539) | | | |
| *Pparg* | CTGTTTTATGCTGTTATGGGTGA | GCTGATTCCGAAGTTGGTGG | 14 | 153 |
| *Runx2* | TCCCTGAACTCTGCACCAAG | GTGGTGGAGTGGATGGATGG | 60 | 152 |
| *U6* | (Predesigned TaqMan assay used; catalogue number 001973) | | | |

**b**

| Gene | Forward primer (5' → 3') | Reverse primer (5' → 3') | Amplicon size (bp) |
|---|---|---|---|
| *App* | GCTCGCTAGCCTCGAATTCTTG TGGTTTGTGGCCC | CGACTCTAGACTCGAAGAAATC AATGTGTATCCTC | 287 |
| *Bcl2* | GCTCGCTAGCCTCGAGCCACAA GTGCCTGCTTTAT | CGACTCTAGACTCGACATCAGC CACGCCTAAAAGT | 4012 |
| *Ccne1* | GCTCGCTAGCCTCGAAAGGAG GGTGCTACTTGACC | CGACTCTAGACTCGATGTTGGC TGACAGTGGAGAA | 233 |
| *Cxcl12*-α | GCTCGCTAGCCTCGACCGAGG AAGGCTGACATCC | CGACTCTAGACTCGAGCCGGAT CTTGTGTTGACTC | 525 |
| *Cxcl12*-β | GCTCGCTAGCCTCGAGCAAGG AAGTGCAGC | CGACTCTAGACTCGACTGCATA TAGGAAGC | 545 |
| *Cxcl12*-γ | GCTCGCTAGCCTCGATAAAACG CTTCTGGAGGCCA | CGACTCTAGACTCGATGAACCC ATCGCTGCTTAGA | 3558 |
| *Cxcl12*-γ (unique part) | GCTCGCTAGCCTCGATAAAACG CTTCTGGAGGCCA | CGACTCTAGACTCGACAAGGA GCACATGACAAGGC | 704 |
| *Icam1* | GCTCGCTAGCCTCGAAGCATTT ACCCTCAGCCACT | CGACTCTAGACTCGAAACCACT GCCAGTCCACATA | 560 |
| *Ing1* | GCTCGCTAGCCTCGACATGTCA GCGAGTGTCAGAC | CGACTCTAGACTCGAGCTGCAC TTCCGGTTATGAT | 490 |
| *Klf7* | GCTCGCTAGCCTCGAGCGGCTA CTCTACTGTCCTT | CGACTCTAGACTCGAAGCATTT GTGACAAAGCGGT | 985 |
| *Pdgfa* | GCTCGCTAGCCTCGAACCAAAC CACACCCAACAAC | CGACTCTAGACTCGAACGTTAA AGGGGCAGAGGAA | 612 |
| *Pin1* | GCTCGCTAGCCTCGACCCTTCC TGCTACTGTCACA | CGACTCTAGACTCGAGCTCTTG GTGTTGCAAGCTA | 1514 |
| *Ptgs1* | GCTCGCTAGCCTCGAAGTGGTT TTGTCTGCCTCCT | CGACTCTAGACTCGAGTCCATC TGTTCCCTCCACA | 636 |
| *Runx2* | GCTCGCTAGCCTCGAATGGCGT CAAACAGCCTCTT | CGACTCTAGACTCGAGACGAC GACAGACTGCTCTA | 636 |
| *Top1* | GCTCGCTAGCCTCGAACAGTGT GGTTTGGGGAAGA | CGACTCTAGACTCGAATGTGGG AATGGACTCTGCA | 1121 |

**c**

| gBlock name/description | gBlock sequence |
|---|---|
| Fragment inserted to create pCMV-ENTRY-neg | TAGGGCGGCCGGGAATTCGTCGACTGGATCGATCCGGTACCGAGGAGATCTtctaga GCCGCCgctagcAAGcatatgACGAGACGCGTACGCGGCCGCTCGAGCAGAAACTCAT CTCAGAAGAGGATC (additional restriction sites shown in lower case) |
| *App* mut | GCTCGCTAGCCTCGAATTCTTGTGGTTTGTGGCCCGGAAAAAACTCCTACTTGAAAT ATGCTTTAAAAATCGATGGGG**C**A**G**GCTTCTTGTGAACGTGGGCGTCTAGCTGCTTCT CCTACGTATTCTTTTCCTGATCACTATGCATTTTGAACATTTTTTTAAGTATTCCAAAT GACTTAGAAAATTCTTTTTCCATGACTGCATCTTACTGTACAGATTGCTGCTTCTGCT CTCTTTGTGATATAGGAATAAGAGGATACACATTGATTTCTTCGAGTCTAGAGTCG |

| | |
|---|---|
| *Cxcl12*-γ (unique) mut | GCTCGCTAGCCTCGATAAAACGCTTCTGGAGGCCAGATCTGTGCTCAAGCCATAGTT<br>CTGCTTAGAAAGG**C**A**G**GCCCCACCCTTACCGGACACTGGGAAGAACTGTTGGCCCC<br>TAGAAACCAAAGGCCAAACTGAGGCTGCCCTGAGTTGGAAGACCACTTTCTGAAAT<br>GCCCATGGACTCTGCCTCCCAACCATTCGTCTCTCACTCCTAGCAGAGCTGTCTGTGC<br>AGACTGTTTCTTAGGAGGCACAGCAAGCTCCAGGGAACCCTCTGTGCTTATGAAGC<br>TCGTCTGGTGGGCAACCCCAGCCCACTGGACAGAGTCCTCATGGAAATGCCTGGGA<br>AGCTGATTTCATCTAAGGATGGGTTGAAGTAGGATGTGCTCCTGCGACTTCTCAGG<br>CAGGTGAGAGGGGTAGTCCTTACACTGTCTAGCATAAACGCCTTCCGGAAGGACCT<br>GCAGCTCCAGAGACCACCTCCTGAGCACCAAGACCTCTTCTGGTGGTGTGGAACCA<br>GCCAAGAGATTTCAAGGAAGAGTGATTATTTGATGAATGCTATGGGAATGGCCTCT<br>TCTCTTGGAGTTCTGAGGCCTGGG**C**A**G**GCCCAGGAACACTGGGCACCTGCTGCTGT<br>TAGGGCCAATGCATAGTCTCAGCACCGGTGTCCTAAGGTTAAGG**A**G**C**TGCGCCTTG<br>TCATGTGCTCCTTGTCGAGTCTAGAGTCG |
| *Pdgfa* mut | GCTCGCTAGCCTCGAACCAAACCACACCCAACAACAAAAAAAAACCCCACCAGAAG<br>AAACAAAAAGACTGAAGTGGGCGACGGAAGGGCCGCCCTGCAGGACTGACCCTCG<br>AGAGGTGGCTGATGTTGGCCACCACCCTGGATTCTGCCCGTGGGCACAGGTGCTTT<br>TCGCCAAGGACGCCTGGAGCCCGCCTGG**C**A**G**GCTGCCTGCCAGCTCCGTGGGAACT<br>GAACAGGTGGGAGATGCCCACGGCCTGGAAGCCACCACCTCGCTTGAGTCTGCCTA<br>GGACCTTCCTGATGGCCTTACGTGGAATTTCTTTATTTTCGTTTTCCTTTCCTTCTTTC<br>TCTCTCTTTTTTACACCATGAAGTGAGTAAGCTTCTCCTTTTTATTCTCTGCCTTCGA<br>GTCTAGAGTCG |
| *Pin1* mut | GCTCGCTAGCCTCGACCCTTCCTGCTACTGTCACACAGTATTTATTGTTCCTAAAATG<br>ACTGGGAGGGGCTCTGAGCATCCCGCTCCCTGTTTGCCCCTATTCGGGGCTGTCCTA<br>GCCAGGCTCCTTCTGGAGGAATTGACTTCAGAAGGG**A**A**C**TGGGGATGGGGAGGCC<br>CCCAGGCTCAGGGGGTCAGAGGTGACCTGATCCCCCAGGTCTCAGAGGCAGGCAG<br>GAGGGCCTTCTTTCTTGTTAGTTATATGCTCCATTCGTTCCTCTGTTCAGTTGCAAAA<br>AGCAGACGCTCCATACCTGGCGGTGCAGCAACTCAGCCTCAGGGTGC**T**G**G**GCAGC<br>ACCCTACGCACCTTCCATTAAATCTGGAACCACTGGTTGGTGGTCCTGTATCATTACC<br>TTCCTGGTGAGGTGTGGGGCCCTTGTCTCAGGGATGGGGCTTTTACCCACGTGTCC<br>AGACTCTTCTCTAGCTCTAAAAGACCTTCTTTTCCACAGCTTGGGGGGTGCTGGCAA<br>GAATTCTTTCCACTAGCTATATTCCCAGGGCTGCAAATACCTCTGAGCCCCACCAGT<br>CCAGACCCACTTGTATAAGCACCCTGGGGACTGTTCAGGTATGCGCAGACCACCCC<br>ACCACCCCACCCCACTTCCAGCTGTTAGGTCTGGAGTGATCCCAAGAGAACCTGGCT<br>TAGCATCTGCCTTGGCACACCCCCGGTGCCCATCCCTCCCCCTCAGCACACAGACAG<br>ACTGCTTATTGGTGAGAACAAATCACCCAGCATCTGCCACCCACTTCTGACTGGATA<br>TTCCTGGGCTCAGCCACAAGACCTAGCACTCTCCAGGTGCTGTTTCCATGGTAACAG<br>AGGCCCCCTCAACTGGAGGAGAGGGGGGCTTTATCAGCTCCCAGAAGAAAAGAGA<br>TCCTATTGGAGGGGGCTGGGAAATGGATATTGCTTTGAGTGGCAGGTGGGGAGAG<br>AAGGCGGACTCTCCCACCTTGGCTAGGGCCCTATGTCTCCCACAACTGCCTCTGAGC<br>CTAAGGACTGAGGGCTCAGAATTCAATCAAGGTTTCTGCATATACCCACAAGTGAT<br>GGGGGGCCCAGCCTGGGGGATGTTCTAACCAGCAAGGCTTGGGGGCCAGAGCAA<br>CATTAAGCCTGGACCCCAAGAGAGCCCTCCCTCCCCACAGGCCTGAGTCTCATGAAC<br>AGAGTCACCTGAGCCTGCTCCCATCTGGCTGCTTTCTAGAGCAATCGCTCTTTGCCA<br>GCCGCCTGCTGGAGCATTTGTCATTAGGAAGGTTTAATGAGTTTTTTCCCAGCTCAG<br>ACAGGCACAGGCTGAGCTGAGTTGGCTATACCTCCAGCAAAATGTTCTCCAGTGGG<br>CCAGTGGGGACTCAAGGCAAACAGTCACAGTTTTGCTGTCTTGCAACAGTTGGGTG<br>GGCTCAGGAGG**A**A**C**TGGCAGCCCTCCAAGTATCATGGGACCTTTGGAGGGAAGAA<br>GAGGACCTCTAGCTTAGCTTGCAACACCAAGAGCTCGAGTCTAGAGTCG |
| *Runx2* mut | GCTCGCTAGCCTCGACTCCAACCTGCGTTTTCCTCAGAATTCATCCTGACTCCCTCTG<br>ACACAGATTGAGGGGGGGGGGAAGAAACCCAACCCGCACCAAAGCAAAACACTTT<br>GCCTTCTAAAGGCTGTGCACCAAGTAGACGCAGATGGTCAGCCCACCTTTGTGTTTC<br>CTTAAGATGGAAATTGTAACTGATGCTATTTATTGTTTGTGTGTGGTAGCTTGAAGC<br>ACACCACGGTCCATGTGTTTGTCTGATACCTATTTCAGCAGCATTTACAAAAGCCAG<br>TGTCTGGTTACACTTTTCAGTTTTCATTAATCAACATGAAAATGTTACCATTGGTGCC<br>AGCTGCACCCTATTTAATTTTTTTAAGGGCACTATATTTGTACATTTCGTTTTTAATGT<br>TAAAGGGCTTCTTAAAGTTTACAGTACAGTTATCAAGGGAATAGAGGG**C**A**G**GCATT |

| | |
|---|---|
| | AGTGCCTAAATGTTATTCTAGTGGCTGCAGGCAGCAACCCAGAAGCAGTTTTGAAA ACAGGTTGTTTCCCTCTGTCCTCCCTTATTTGGGAAAATTCAAGTGCTTTCTTCACCTT TCAGGCACCTCACGGTGACTCCCGTTACTTAGAGCAGTCTGTCGTCGTCTCGAGTCT AGAGTCG |

**Supplementary Table S3 - Oligonucleotides used in this research project.** (**a**) Primers and Universal Probe Library probes used to assess gene expression using RT-qPCR. (**b**) PCR primers used to amplify 3'UTR fragments of putative miR-324 target genes. (**c**) gBlocks utilised for molecular cloning in this project. All sequences in this table are shown in a 5' to 3' orientation. For mutant (mut) sequences, substituted nucleotides are shown in bold and underlined.

| **Protein** | **Dilution/concentration** | **Host animal** | **Purchased from** |
|---|---|---|---|
| β-tubulin | 1:1000 | Mouse | Sigma-Aldrich, catalogue number T8328 |
| FLAG (Myc-DDK) | 1:2000 | Mouse | Cell Signalling Technology, catalogue number 2276S |
| Runx2 | 1:500 | Rabbit | Cell Signalling Technology, catalogue number 125 |

**Supplementary Table S4 - Primary antibodies used for western blotting.**
